# Scrib organizes cortical actomyosin clusters to maintain adherens junctions and angiogenic sprouting

**DOI:** 10.1101/2025.04.01.645927

**Authors:** Lakyn N. Mayo, Fiona Duong, Ana Mompeón, Kyle A. Jacobs, M. Luisa Iruela-Arispe, Matthew L. Kutys

## Abstract

Spatiotemporal control of adherens junction fluidity and integrity is critical for angiogenesis, but underlying mechanisms are incompletely understood. To identify unappreciated regulators of endothelial adherens junctions, we performed VE-cadherin proximity ligation mass spectrometry, revealing significant interaction with the multifunctional scaffold Scrib. Utilizing a 3D angiogenesis-on-chip model, we find Scrib-depleted microvessels generate reduced intact sprouts and increased single-cell detachments. This defect was characterized by adherens junction instability and decreased actomyosin in the junctional cortex, yet was not caused by changes in catenin-dependent VE-cadherin coupling to actin. Instead, Scrib controls the formation of cortical actomyosin clusters, which critically organize the architecture and dynamics of the junctional actomyosin cortex to promote adherens junction stability. We further discovered that unconventional myosin-1c is a critical effector linking Scrib cortical dynamics and VE-cadherin to stabilize adherens junctions during angiogenic initiation. Our results demonstrate a new role for Scrib directing cortical actomyosin organization that is critical for precise control of adherens junctions during angiogenesis.

## INTRODUCTION

Angiogenesis, the growth of new blood vessels from previously existing vessels, is essential for tissue growth and injury repair (Herbert and Stainier, 2011). Despite extensive research, we still lack complete understanding of how angiogenesis is spatially and temporally regulated to drive proper neovascularization therapeutically in settings such as tissue ischemia (Kanazawa et al., 2019) and tissue-engineered organs (Cancedda et al., 2020), or to block vascularization of tumors (Martin et al., 2019). A key unmet need is a deeper understanding of angiogenic initiation, whereby angiogenic stimuli trigger the basal invasion of a tip cell from a uniform polarized monolayer of endothelial cells (ECs) (Guo et al., 2024). A core aspect of this event is controlled remodeling of tip cell cell-cell adhesions, which permits a tip cell to elongate and migrate without detaching from the parent vessel and later, stalk cells which follow (Cao et al., 2017).

Adherens junctions (AJs) mediate cell-cell adhesion and serve key chemo-mechanical signaling roles to regulate stability and morphogenesis of multicellular tissues (Capuana et al., 2020; Mège and Ishiyama, 2017). AJ destabilization through remodeling of VE-cadherin, the primary cell-cell adhesion receptor in endothelial AJs, is crucial for angiogenesis (Abraham et al., 2009; Cao et al., 2017; Grimsley-Myers et al., 2020; Lampugnani et al., 2003; Szymborska and Gerhardt, 2018). This is mediated by several processes, primarily endocytosis and recycling of VE-cadherin (Grimsley-Myers et al., 2020; Malinova et al., 2021) and internalization and morphologic changes via activity of junction-proximal cortical actomyosin (Abraham et al., 2009; Cao et al., 2017), hereafter referred to as junctional actomyosin. As sprouting continues, AJ stability is tuned between neighboring cells to permit cell rearrangements within the sprout (Bentley et al., 2014). This delicate, dynamic control of AJ integrity is essential for multicellular sprouting, yet underlying control systems still remain to be characterized.

VE-cadherin assembles multimeric AJ plaques that include p120 catenin, which shields VE-cadherin from endocytosis (Nanes et al., 2012), and β-catenin and ⍺-catenin, which mechanically link VE-cadherin to the underlying cortical actomyosin network (Cao and Schnittler, 2019; Efimova and Svitkina, 2018; Lampugnani et al., 1995). Organization and contractility of the cortical actomyosin network is central to AJ stability and remodeling (Mège and Ishiyama, 2017). Indeed, efficient angiogenesis and AJ fluidity are coupled to the regulation of cortical actomyosin, including reduced contractility at AJs and formation of junction-associated lamellipodia (Abraham et al., 2009; Cao et al., 2017). Despite this destabilization, mechanical integrity of the AJ must be maintained so that AJs elongate and cell-cell adhesion is not compromised (Sauteur et al., 2014; Wimmer et al., 2012). Vinculin, a mechanosensitive protein that reinforces the interaction between ⍺-catenin and actin, prevents endothelial AJs from breaking under mechanical forces, crucial for sufficient angiogenic sprout density and nascent lumen formation (Carvalho et al., 2019; Huveneers et al., 2012; Kotini et al., 2022). Raf-1-driven Rho-kinase activity at AJs is required to maintain cohesion in nascent sprouts (Wimmer et al., 2012), and formin-polymerized actin cables at AJs are required for nascent lumen formation and sprout continuity (Phng et al., 2015). In other systems, supramolecular cortical actomyosin architectures, including clusters/nodes (Chou et al., 2024; Wollrab et al., 2016) and sarcomere-like arrays (Yu-Kemp et al., 2021), have been increasingly recognized to promote AJ remodeling during tissue morphogenesis (Kruse et al., 2024; Rauzi et al., 2010). Altogether, while precise remodeling of both AJs and cortical actomyosin are crucial for angiogenesis, we lack a complete mechanistic understanding of this reciprocity and associated molecular regulators.

To identify unappreciated regulators of endothelial AJ stability, we profiled VE-cadherin interactors using unbiased proximity biotinylation (BioID) and mass spectrometry. Among several new interactors identified was the multifunctional scaffold Scrib, a central orchestrator of diverse forms of tissue morphogenesis (Bonello and Peifer, 2019). Here, employing 3D engineered models of primary human vasculature wherein we investigate mechanisms underlying angiogenic initiation at high spatiotemporal resolution, we find that Scrib deletion (*SCRIB^KO^)* impairs angiogenic sprout integrity by increasing tip cell detachment from the tip cell-to-parent vessel interface. The *SCRIB^KO^*defect arises from disrupted AJs and junctional actomyosin but surprisingly occurs independently of cadherin-catenin interactions. Instead, we find that Scrib directly regulates cortical actomyosin architecture through organizing clusters in the junctional cortex, which promotes AJ stability and maintains endothelial mechanics. Further, we demonstrate that Scrib regulates tip cell integrity and VE-cadherin stability through a novel association with unconventional myosin-1c (Myo1c). Altogether, we define a new mechanism by which Scrib directs cortical actomyosin organization to stabilize endothelial AJs that is necessary to maintain multicellular integrity during vascular sprouting.

## RESULTS

### VE-cadherin proximity ligation mass spectrometry reveals interaction with Scrib

To identify unknown regulators of VE-cadherin, we performed unbiased proximity ligation mass spectrometry using BioID (Kim et al., 2014). VE-cadherin was tagged at the C-terminus with the promiscuous biotin ligase BirA* (VE-cadherin-BioID, VE-BirA*) and an HA-tag and stably expressed in primary human microvascular endothelial cells (hMVECs) (Fig. 1a-i). VE-cadherin-BioID expressing monolayers were incubated with media containing biotin overnight, biotinylated interacting proteins were isolated via streptavidin affinity purification (Fig. 1a-ii), and then analyzed by mass spectrometry (Fig. 1a-iii). hMVECs expressing a cytoplasmic BirA* (Cyto-BirA*) were used as control. Analysis of the most abundant VE-cadherin-BioID interactors expectedly revealed core AJ proteins that associate with the intracellular domain of VE-cadherin (Fig. 1b), including ⍺-catenin, β-catenin, junction plakoglobin, and p120 catenin (Cao and Schnittler, 2019; Efimova and Svitkina, 2018; Grimsley-Myers et al., 2020; Lampugnani et al., 1995). However, several unappreciated proteins were identified to interact with VE-cadherin, including Scribble (Scrib), which we subsequently validated via streptavidin pull-down and Western blot of VE-cadherin-BioID biotinylated proteins (Fig. 1c). Scrib is a multifunctional adaptor that localizes to the basolateral plasma membrane of epithelial cells and plays important roles in epithelial polarity and cell-cell adhesion (Awadia et al., 2019; Choi et al., 2019; Hendrick et al., 2016; Lohia et al., 2012; Qin et al., 2005). Despite the substantial structural and polarity differences that exist between epithelial and endothelial cells (Lee and Bautch, 2011; Lizama and Zovein, 2013; Worzfeld and Schwaninger, 2016), a role for Scrib at vascular AJs has not been investigated.

**Figure 1:**
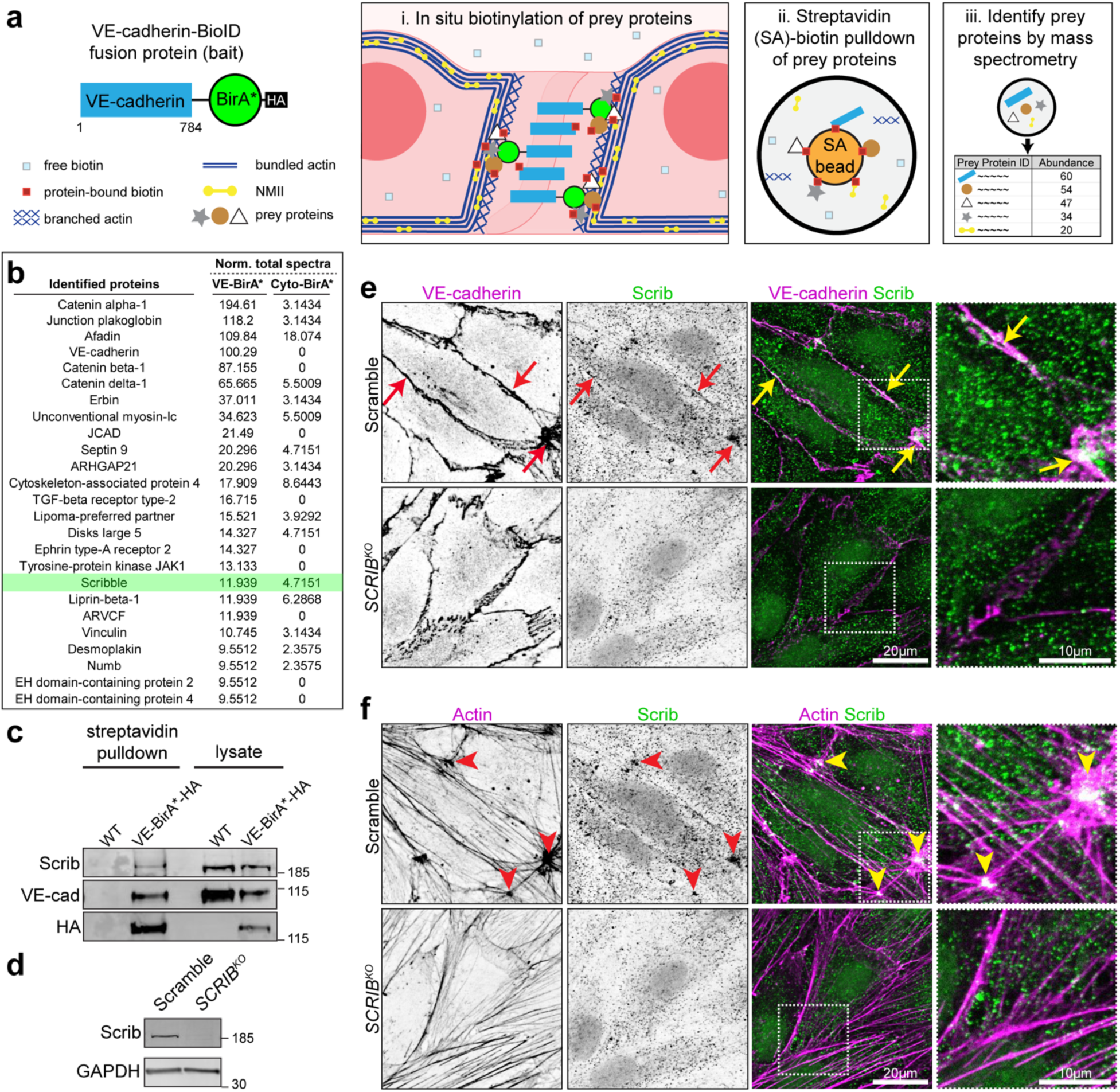
VE-cadherin proximity ligation mass spectrometry reveals interaction with Scrib. **(a)** Proximity ligation mass spectrometry using VE-cadherin-BioID to identify unappreciated proteins in proximity to the intracellular domain of VE-cadherin. BirA*, a promiscuous biotin ligase, was fused to the intracellular domain of VE-cadherin as a “bait” fusion protein construct. **(i)** VE-cadherin-BirA* was introduced into wild-type primary human microvascular endothelial cells (hMVECs) via lentiviral transduction. Free biotin was introduced into cells, and VE-cadherin-BirA* biotinylated proximal “prey” proteins. **(ii)** hMVEC lysates were incubated with streptavidin beads to extract (pulldown) biotinylated proteins. **(iii)** Biotinylated proteins were eluted from the streptavidin beads and analyzed via mass spectrometry. **(b)** Biotinylated proteins identified by mass spectrometry that are selectively enriched by VE-cadherin (VE-cad-BirA*) compared to a cytoplasmic control (Cyto-BirA*) are listed according to abundance. **(c)** Western blot of affinity-purified biotinylated proteins from wild-type or VE-cad-BirA* expressing hMVECs immunostained for Scrib, VE-cadherin, and HA. HA immunolabels VE-cadherin-BirA*. **(d)** Western blot of Scramble and *SCRIB^KO^* lysates immunoblotted for Scrib and GAPDH. **(e)** Scramble and *SCRIB^KO^*fluorescence micrographs of Scrib and VE-cadherin (colocalization, arrows) and **(f)** Scrib and cortical actin clusters (colocalization, arrowheads) the same field of view. (N = 3 independent experiments).

Endothelial-specific *Scrib* deletion causes barrier function defects and increased risk of atherosclerosis in mice (Schürmann et al., 2019); however, if and how Scrib influences endothelial polarity, cell-cell adhesion, and vascular morphogenesis remains unknown. To assess Scrib localization and function in hMVECs, we applied our previously established CRISPR-Cas9 methods to generate scrambled sgRNA (Scramble) control and Scrib knockout (*SCRIB^KO^*) hMVECs (Polacheck et al., 2017; Ran et al., 2013; White et al., 2023) (Fig. 1d). We first assessed Scrib localization by immunofluorescence and indeed observed that a subset of Scrib co-localizes with VE-cadherin at endothelial AJs (Fig. 1e, arrows). Curiously, we also observed that a larger subset of Scrib is organized into clusters that strikingly overlap nodes of interacting actin fibers within the cell cortex (Fig. 1f, arrowheads), and these Scrib clusters occasionally overlap both actin nodes and VE-cadherin. Scrib immunostaining at either localization was abolished in *SCRIB^KO^* cells. Scrib localization was not altered upon knockout of Erbin or Disks large 5 (DLG5), regulators of epithelial cell polarity (Choi et al., 2019; Liu et al., 2017) that appeared in the VE-cadherin-BioID screen (Fig. 1b, Fig. S1a, b). Altogether, our data indicate that Scrib is a VE-cadherin interacting protein that localizes both to AJs as well as cortical actin nodes in primary human endothelial cells.

### Scrib maintains nascent sprout integrity during angiogenesis

Global *Scrib* disruption in animals causes vascular defects, and Scrib-depleted endothelium in vitro exhibits disrupted directed migration, suggesting a role for Scrib in vascular morphogenesis (Michaelis et al., 2013). However, a molecular, endothelial-intrinsic role of Scrib contributing to angiogenesis remains unclear. To investigate how Scrib might influence angiogenesis, we adapted a previously established microphysiological system that recapitulates and permits the visualization of angiogenesis from human microvessels with high spatiotemporal resolution (Nguyen et al., 2013; Wang et al., 2020). In this model, a 3D microvessel cultured under fluid flow is exposed to a gradient of angiogenic chemokine. To confirm that this platform enables robust assessment of angiogenic sprout formation in hMVECs, we introduced a well-established endothelial chemoattractant, sphingosine-1-phosphate (S1P) (Paik et al., 2001; Wang et al., 2020), to the chemokine channel. Indeed, exposure to S1P gradients initiates reproducible sprouting events from the parent microvessel towards the angiogenic gradient within 24h (Fig. 2a-c, Fig. S2a).

**Figure 2:**
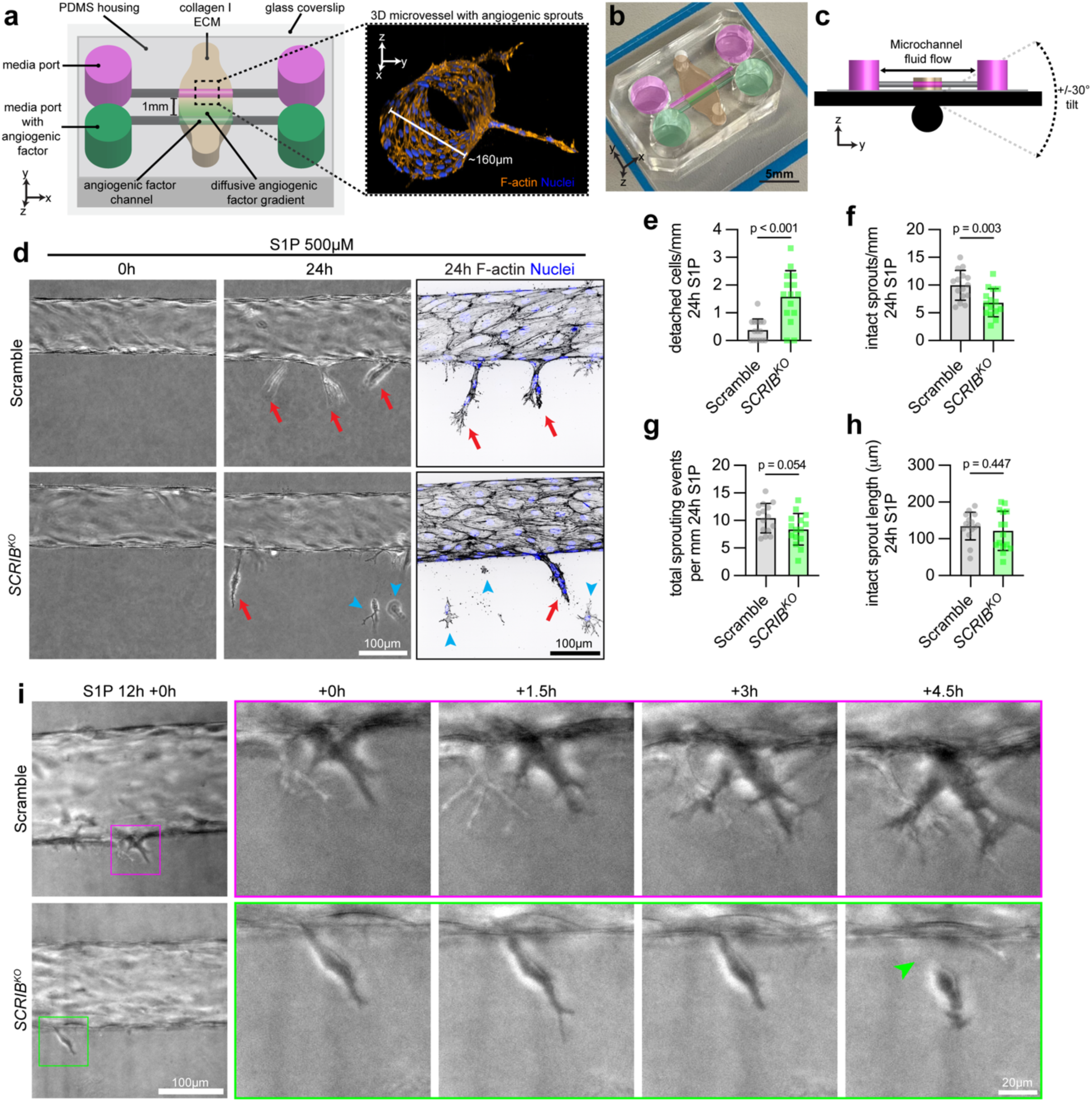
Scrib maintains nascent sprout integrity during angiogenesis. **(a-c)** 3D microfluidic angiogenesis model. **(a)** Key components highlighted on a diagram of the device. Two 3D microchannels of 160 μm diameter are positioned 1 mm apart in a collagen I matrix. hMVECs are seeded onto the top channel (pink), and both channels are supplied by large media ports. After microvessel maturation in the top microchannel, angiogenic factors are flowed through the bottom microchannel (green), generating a diffusive gradient of angiogenic factors such as S1P. The inset depicts an example 3D microvessel generated within the device with directional sprouts towards an angiogenic gradient. **(b)** Photograph of the microfluidic device, pseudo-colored with key components in (a). **(c)** Diagram depicting the mechanism by which flow is applied to the microvessel. **(d)** Representative time course of microvessel angiogenesis stimulated by 500 μM S1P in Scramble and *SCRIB^KO^*, visualized by both phase and confocal microscopy. Red arrows indicate intact sprouts; blue arrowheads indicate detached single cells. **(e-h)** Quantification of sprouting event parameters after 24 h of S1P stimulation (N = 3 independent experiments, n = 3-5 devices per condition per N, unpaired two-tailed t-tests for e-h). **(e)** Number of detached cells per 1 mm parent vessel. **(f)** Number of intact sprouts per 1 mm parent vessel. **(g)** Total number of sprouting events, calculated by number of multicellular sprouts + number of detached cells, per 1mm parent vessel. **(h)** Length of longest intact sprout in each device. **(i)** Representative live imaging time course of a sprout emerging from the parent vessel for Scramble and *SCRIB^KO^*, starting 12 h post S1P initiation (N = 3 independent experiments, n = 1 device per condition per N) (Video 1).

In control microvessels composed of Scramble hMVECs, many intact, multicellular sprouts form over 24h S1P treatment (Fig. 2d, e; Fig. S2a). *SCRIB^KO^*hMVECs form intact microvessels comparably to Scramble cells (Fig. S2b). Yet, in response to the same S1P gradient, *SCRIB^KO^* microvessels form significantly fewer intact, multicellular sprouts, and exhibit significantly more detached single tip cells from the parent vessel (Fig. 2d-f, Fig. S1b). Notably, total sprouting events, the sum of intact sprouts and detached cells, is not significantly different between Scramble and *SCRIB^KO^*(Fig. 2g), suggesting that the overall capacity to initiate a sprout is unchanged between Scramble and *SCRIB^KO^* microvessels. In the few intact, multicellular *SCRIB^KO^* sprouts, there was no significant difference in overall sprout length between Scramble and *SCRIB^KO^* (Fig. 2h), indicating that the detached cell events are likely primarily failures of maintaining cell adhesion upon tip cell emergence. To test this directly, we performed live imaging of sprouting microvessels (Fig. 2i, Video 1). In control microvessels, Scramble tip cells elongate and maintain intact connections with the parent vessel over time. In contrast, *SCRIB^KO^* tip cells elongate, but eventually detach from the parent vessel monolayer. Altogether, we demonstrate that *SCRIB^KO^* impairs angiogenesis in a biomimetic human microphysiological system and specifically identify that Scrib regulates nascent sprout integrity by maintaining tip cell-to-parent microvessel adhesions.

### Scrib regulates endothelial adherens junction linearity independent of cadherin-catenin interactions

AJ remodeling is required for angiogenic initiation, and *SCRIB^KO^*causes tip cell detachments. Scrib impacts epithelial morphogenesis, often through action at AJs (Awadia et al., 2019; Hendrick et al., 2016). We therefore hypothesized that *SCRIB^KO^*may disrupt AJ integrity leading to tip cell detachments. To test, we performed immunofluorescence staining of VE-cadherin and prominent AJ regulators of VE-cadherin stability: ⍺-catenin, β-catenin, and p120 catenin (Cao and Schnittler, 2019; Dartsch et al., 2014; Grimsley-Myers et al., 2020), and quantified AJ linearity and width (Brezovjakova et al., 2019). As a positive control for disrupting AJs, we compared *SCRIB^KO^* to *CTNNA1^KO^*, a CRISPR-Cas9 mediated knockout of ⍺-catenin (Fig. S3a), which is established to disrupt AJ stability (Hayer et al., 2016; Livshits et al., 2012; Noordstra et al., 2023; Zhu et al., 2021). In *CTNNA1^KO^*, the junctional fluorescence intensities of VE-cadherin, ⍺-catenin, β-catenin, and p120 catenin are significantly decreased (Fig. 3a-e). Junctional fluorescence intensities of VE-cadherin and p120 catenin are slightly decreased in *SCRIB^KO^*hMVECs (Fig. 3a-e), while the total protein level of VE-cadherin and p120 catenin are unchanged (Fig. 2f). There was also no change in N-cadherin fluorescence intensity in *SCRIB^KO^* relative to Scramble (Fig. S3b). However, in contrast to *CTNNA1^KO^,* co-immunoprecipitation of VE-cadherin with β-catenin or p120 catenin is not perturbed by *SCRIB^KO^* (Fig. 2g), and endocytic internalization of VE-cadherin in *SCRIB^KO^* is not different from Scramble (Fig. S3c, d).

**Figure 3:**
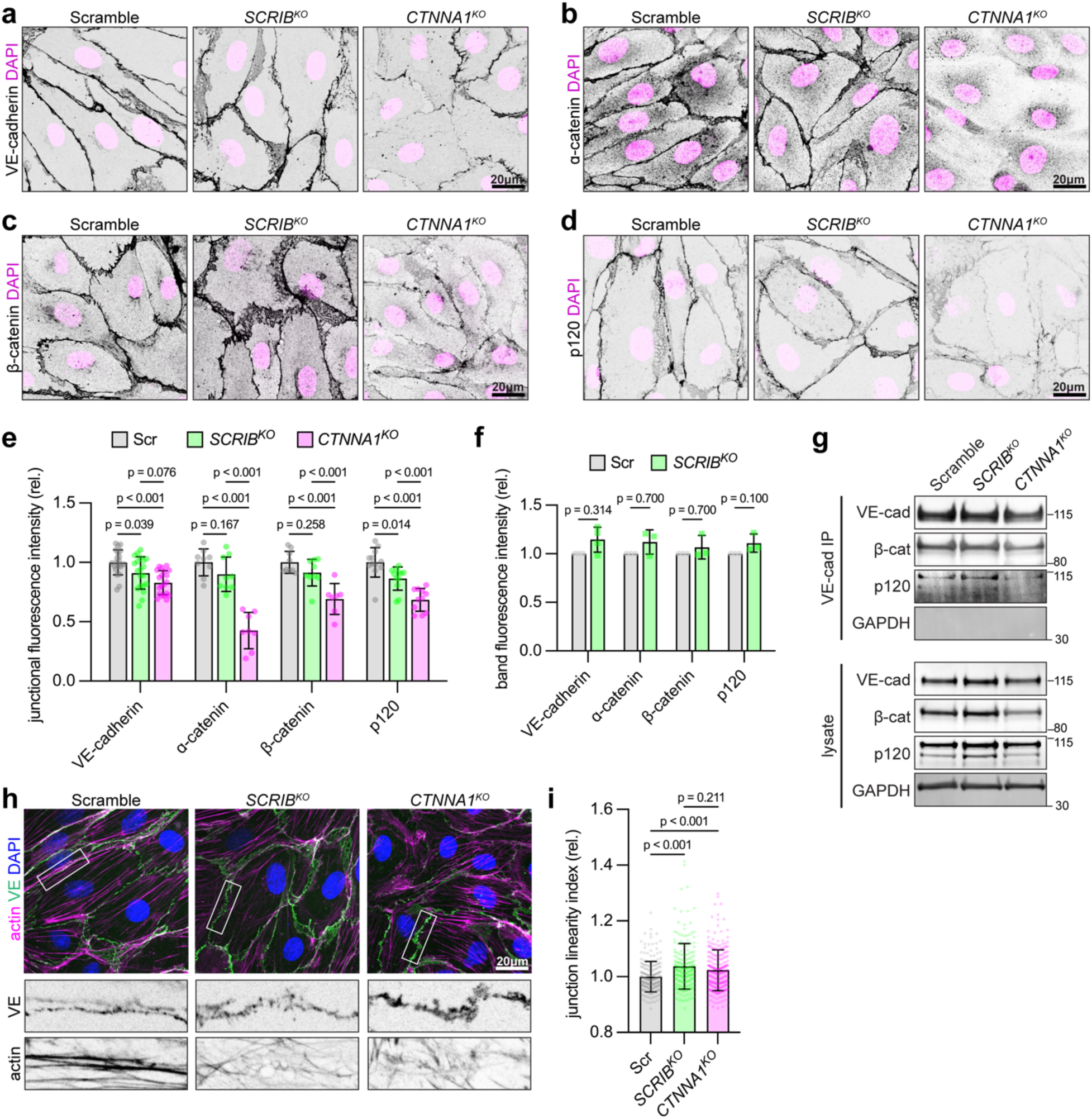
Scrib regulates endothelial adherens junction linearity independent of cadherin-catenin interactions. **(a-d)** Immunofluorescence micrographs of AJ components in Scramble, *SCRIB^KO^*, and *CTNNA1^KO^* hMVECs. **(e)** Quantification of VE-cadherin and catenin junctional fluorescence intensities in Scramble, *SCRIB^KO^* and *CTNNA1^KO^* hMVECs (N ≥ 3 independent experiments, n ≥ 2 patches of 100μm^2^ per condition per N, two-way ANOVA with Tukey’s multiple comparisons test). **(f)** Relative protein expression in Scramble and *SCRIB^KO^* lysates quantified from Western blot normalized to GAPDH. **(g)** Western blot of immunoprecipitation of VE-cadherin from Scramble and *SCRIB^KO^* lysates, blotted for VE-cadherin (N = 3), β-catenin (N = 3), and p120 catenin (N = 2). **(h)** Immunofluorescence micrograph of VE-cadherin, actin, and DNA in Scramble, *SCRIB^KO^*, and *CTNNA1^KO^*hMVECs. **(i)** Quantification of junctional linearity index, defined as the junction contour length divided by the straight-line contour length. (N = 4 independent experiments, n ≥ 40 junctions per condition per N, Mann-Whitney test). Corresponding junctional actin displayed below each representative junction.

Despite key differences between *SCRIB^KO^* and *CTNNA1^KO^*AJ composition, we observed similarities in VE-cadherin phenotype. While junction width was not significantly different in *SCRIB^KO^* or *CTNNA1^KO^*from control (Fig. S3e), we found that *SCRIB^KO^* and *CTNNA1^KO^*both robustly exhibit an increase in junction undulation (Fig. 3h, i), quantified as the linearity index, which is the ratio of the adhesion contour length to the adhesion straight line length. A linearity index greater than 1 indicates junction is more undulated, an indicator of AJ instability (Brezovjakova et al., 2019; Kim and Cooper, 2018; Schulte et al., 2011). Junction undulation has also been linked to disruption of junctional actomyosin (Brezovjakova et al., 2019; Otani et al., 2006; Sri-Ranjan et al., 2022), which we indeed observe in associated *SCRIB^KO^* and *CTNNA1^KO^*junctional actin (Fig. 3h). Altogether, we observe that both *SCRIB^KO^* and *CTNNA1^KO^* disrupt VE-cadherin junction morphology, yet, in contrast, the association of key AJ proteins is maintained in *SCRIB^KO^.* Given the prominent changes we observed in associated cortical actin organization, we next investigated whether Scrib directly influences junctional actomyosin.

### Scrib organizes cortical actomyosin through a mechanism distinct from catenin-dependent cadherin coupling to actin

*SCRIB^KO^* disrupts AJ linearity (Fig. 3), which is often a sign of changes in underlying junctional actomyosin (Brezovjakova et al., 2019; Sri-Ranjan et al., 2022). Scrib also exhibits a striking localization to cortical actin clusters (Fig. 1f) in concert with AJs (Fig. 1e), and remodeling of VE-cadherin by cortical actomyosin is important for angiogenic sprouting (Abraham et al., 2009; Cao et al., 2017). Therefore, we asked if Scrib regulates VE-cadherin stability and angiogenic sprout integrity through control of cortical actomyosin.

We examined key components of the actomyosin cortex in hMVECs: non-muscle myosin isoforms IIB and IIA (NMIIB/NMIIA) and phosphorylated myosin light chain (pMLC, phosphorylated at Ser19), via immunofluorescence co-staining with VE-cadherin (Fig. 4a-c). By creating a dilated junctional mask, we quantified the abundance of actomyosin staining within the mask (“cortical”) relative to actomyosin staining outside the mask (cell “interior”) (Fig. 4d). Of the three actomyosin components examined, interestingly NMIIB was the most abundantly localized to the junctional cortex in Scramble (Fig. 4d). In *SCRIB^KO^*, NMIIB, NMIIA, and pMLC intensities are decreased at junctions (Fig 4a-d), corresponding to a significant decrease in cortical:interior ratio for NMIIB, NMIIA, and pMLC in *SCRIB^KO^* to near 1, indicating that the abundance of actomyosin in the junctional cortex is similar to the abundance of actomyosin in the cell interior. Total protein levels of NMIIB, NMIIA, and ppMLC (phosphorylated at Thr18/Ser19) were unchanged in *SCRIB^KO^* (Fig. S3f, g), suggesting this result is due to protein mislocalization. Thus, Scrib maintains junctional localization of NMIIB, NMIIA, and pMLC in the endothelial cell cortex.

**Figure 4:**
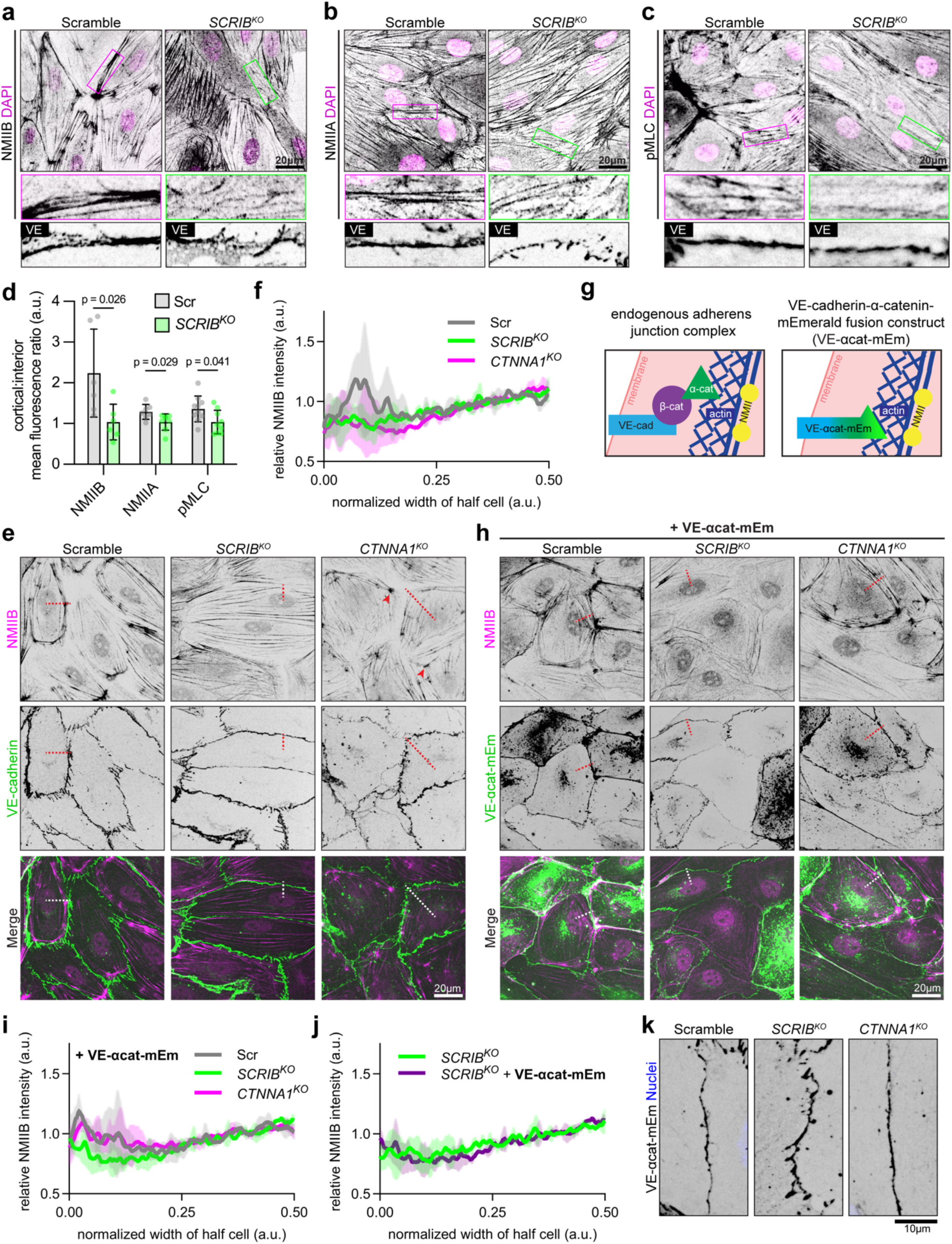
Scrib maintains cortical actomyosin organization in a mechanism distinct from catenin-dependent cadherin coupling to actin. **(a-c)** hMVEC AJ-proximal cortical actomyosin components visualized by immunofluorescence co-stain with VE-cadherin: NMIIB, NMIIA, and pMLC. **(d)** Quantification of relative NMIIB, NMIIA, and pMLC cortical abundance by taking the ratio of cortical mean fluorescence intensity (using a dilated junctional mask) to the mean fluorescence intensity in the rest of the cell (interior) (N = 3 independent experiments, n = 1-4 patches of 100μm^2^ per condition per N, Mann-Whitney test). **(e**) Immunofluorescence micrographs of NMIIB and VE-cadherin in Scramble, *SCRIB^KO^*, and *CTNNA1^KO^* hMVECs. **(f)** NMIIB cortical abundance via immunofluorescence, measured by fluorescence intensity line scans from the junction through the brightest cortical NMIIB to the midpoint of the cell. Fluorescence intensity values were normalized to the non-cortical abundance (latter 50% of line scan) (N = 3 independent experiments, n ≥ 20 cells per condition per N). **(g)** Schematic of VE-cadherin-α-catenin-mEmerald fusion construct (VE-αcat-mEm). **(h) I**mmunofluorescence micrographs of NMIIB and VE-αcat-mEm in Scramble, *SCRIB^KO^*, and *CTNNA1^KO^* hMVECs expressing VE-αcat-mEm. **(i, j)** NMIIB cortical abundance via immunofluorescence in VE-αcat-mEm expressing cells. (N = 3 independent experiments, n ≥ 20 cells per condition per N). **(k)** Representative junctions between cells expressing VE-αcat-mEm.

VE-cadherin connects to cortical actomyosin through interaction with β-catenin which complexes with ⍺-catenin, which binds directly to actin (Duong et al., 2021). While *SCRIB^KO^* does not affect the interaction between VE-cadherin and catenins (Fig. 2g), we posited the disrupted cortical actomyosin organization in *SCRIB^KO^* might be secondary to a failure of catenin-dependent coupling of VE-cadherin to cortical actomyosin. To test this, we first analyzed the distribution of NMIIB, the most cortically enriched actomyosin component we observed in hMVECs (Fig. 4a, d), in *CTNNA1^KO^* compared to *SCRIB^KO^*. As expected, Scramble exhibits peak NMIIB fluorescence in the cell cortex (Fig. 4e, f). Similar to *SCRIB^KO^,* we observed a lack of NMIIB cortical enrichment in the cell cortex in *CTNNA1^KO^* (Fig. 4e, f). Interestingly, we noticed a different pattern of disruption in *SCRIB^KO^*, wherein NMIIB cortical clusters are absent, compared to *CTNNA1^KO^* wherein NMIIB clusters were retained (arrowheads) (Fig. 4e).

To further test whether the *SCRIB^KO^* NMIIB cortical localization defect was a result of disrupted catenin-dependent cadherin coupling to actin, we utilized a previously published chimeric construct fusing VE-cadherin to the C-terminus of ⍺-catenin and fluorescent mEmerald (VE-⍺cat-mEm), which lacks β-catenin binding but allows binding to actin, vinculin, and ⍺-actinin (Fig. 4g) (Dartsch et al., 2014; Imamura et al., 1999; Schulte et al., 2011). When expressed in ECs, this construct increases VE-cadherin association with actin and increases junction linearity (Dartsch et al., 2014; Schulte et al., 2011). As validation, expressing VE-⍺cat-mEm in *CTNNA1^KO^* results in restoration of NMIIB in the junctional cortex to a level near identical to Scramble (Fig. 4h, i, Fig. S3h). However, NMIIB cortical localization is not restored in *SCRIB^KO^* expressing VE-⍺cat-mEm (Fig. 4h-j). Consistent with these observations, we found increased VE-cadherin junction linearity in *CTNNA1^KO^*, but not *SCRIB^KO^*, cells expressing VE-⍺cat-mEm (Fig. 4k). Thus, we conclude that Scrib regulates VE-cadherin by maintaining cortical actomyosin organization through a mechanism separate from catenin-dependent cadherin coupling to cortical actomyosin.

### *SCRIB^KO^* disrupts cortical actin clusters and dynamics

Scrib strikingly localizes to endothelial cortical actin clusters (Fig. 1) and maintains actomyosin abundance in the junctional cortex through a mechanism that does not impact cadherin-catenin interaction (Fig. 4). Actomyosin clusters are meso-scale structures that are increasingly recognized to integrate forces for morphogenetic activities such as apical constriction (Martin et al., 2009), and wound closure (Bhat et al., 2024). Next, we asked whether Scrib controls cortical actomyosin organization through actomyosin dynamics at observed clusters. In Scramble hMVECs, actin is arranged in large bundles in the endothelial cortex, with increased fluorescence intensity in the junctional cortex than in the cell interior (Fig. 5a, b). At the ends of these actin bundles, which circumferentially connect to other bundles at an angle, nodes of clustered actin structures are often present (arrows), and this is precisely where Scrib is enriched (arrows) (Fig. 5a). In *SCRIB^KO^* hMVECs, these clusters are nearly abolished (arrows depict where a cluster would be located in Scramble cells) (Fig. 5a). This loss is accompanied by lacier actin bundles in the cortex, with a lower cortical:interior ratio (Fig. 5a, b), similar to previous observations of NMIIB/A and pMLC (Fig. 4a-d).

**Figure 5:**
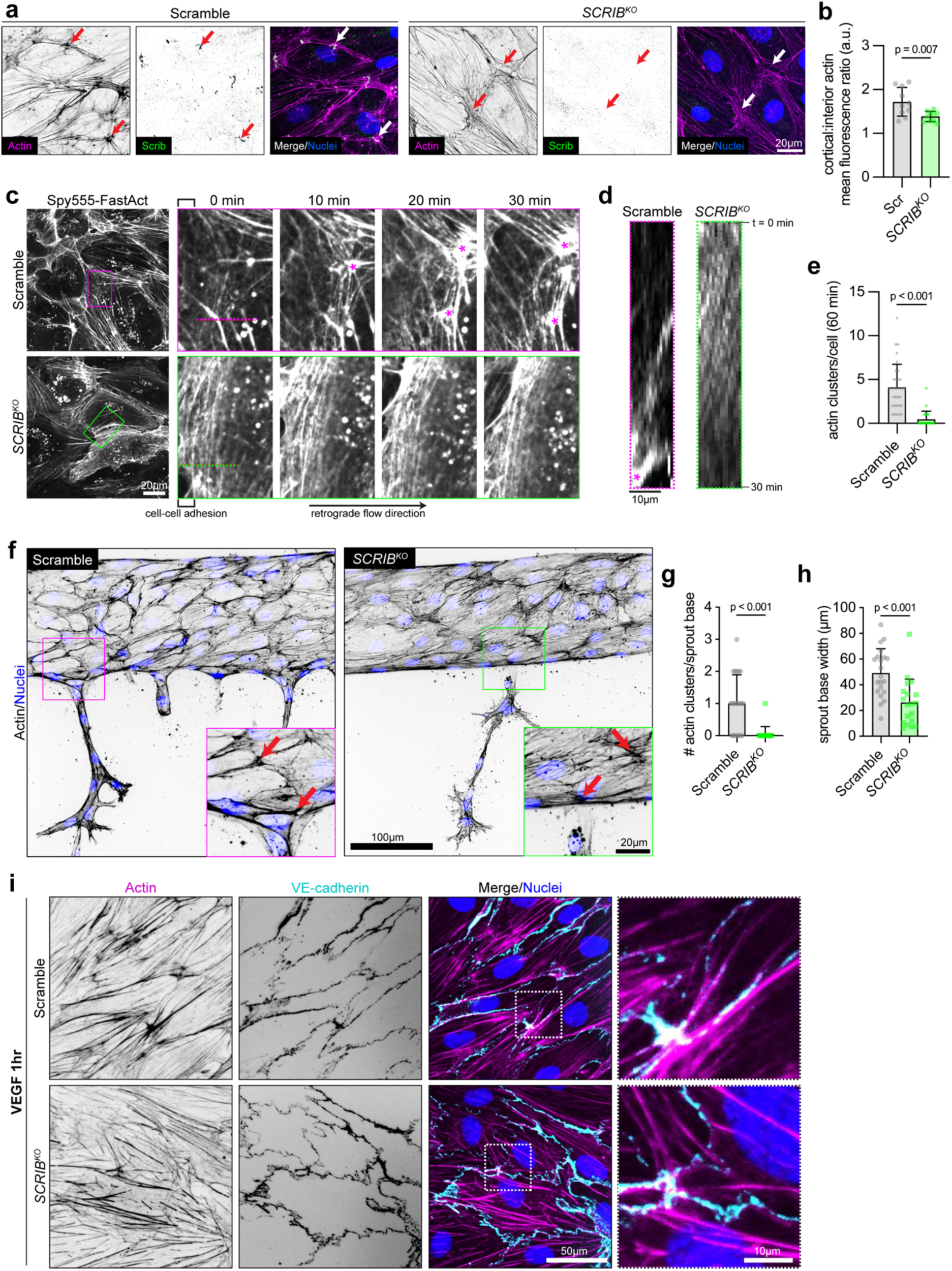
Scrib localizes to cortical actin clusters, which are present at the base of intact sprouts and correlate with linear adhesions upon angiogenic stimulation. **(a)** Scrib immunostaining and actin labeling (overlap at clusters, arrows) in Scramble and *SCRIB^KO^* hMVECs. **(b)** Ratio of cortical mean fluorescence intensity of actin (using a dilated junctional mask) to the mean fluorescence intensity of actomyosin in the rest of the cell (interior) (N = 5 independent experiments, n = 2 patches of 100 μm^2^ per condition per N, Mann-Whitney test). **(c)** Live actin imaging via incubation of Spy555-FastAct, highlighting a representative region of retrograde actin flow from a cell-cell adhesion in the inset (clusters, asterisks) (Video 2). **(d)** Representative kymograph from dotted lines of inset in (b). **(e)** Quantification of actin clusters present per cell during 60 min of live imaging (N = 3 independent experiments, n ≥ 6 cells per condition per N, Mann-Whitney test). **(f)** Representative sprout-parent vessel interfaces in Scramble (actin clusters, arrow), and *SCRIB^KO^*(actin enrichments, arrows). **(g)** Quantification of actin clusters within an ROI of 25 x 50 μm spanning at the sprout-parent vessel interface in Scramble and *SCRIB^KO^* (N = 4 independent experiments, n ≥ 5 sprouts per condition per N, Mann-Whitney test). **(h)** Quantification of sprout base width for the same sprouts quantified in (g). **(i)** Representative images of confluent endothelial monolayers grown on compliant polyacrylamide gels and treated with VEGF (50 ng/ml, 1 h), co-stained for actin and VE-cadherin for Scramble and *SCRIB^KO^* (N = 3 independent experiments).

Timelapse imaging of actin in Scramble hMVECs revealed actin clusters to be dynamic structures that coalesce from retrograde flow of actin fibers from cell-cell junction interfaces in confluent monolayers (Fig. 5c, Video 2). Clusters often merge with other clusters and become larger in size (Fig. S4a, b; Video 3). In contrast, retrograde actin flow from cell-cell interfaces in *SCRIB^KO^*is unable to cluster (Fig. 5c, Video 2). A median of 4 actin clusters, defined as dynamic high-intensity actin structures joining several actin bundles at an angle, were present per cell in Scramble during 60 minutes of live imaging (Fig. 5d), with 100% of cells observed exhibiting at least 1 cluster. Kymograph analysis revealed comparable rates of retrograde actin flow from cell-cell adhesions in both Scramble and *SCRIB^KO^*, however, in *SCRIB^KO^*, individual actin fibers fail to coalesce into actin clusters (Fig. 5e, dashed lines in Fig. 5c). *SCRIB^KO^* cells occasionally form overlapping actin arrays from retrograde flow of actin, but these arrays typically fail to coalesce into clusters (Fig S4c, d; Video 4), a median exhibiting 0 clusters per cell over 60 minutes of live imaging (Fig. 5d). We conclude that organized actin clusters are prevalent and dynamic in the junctional cortex of confluent Scramble endothelium, while they are nearly abolished in *SCRIB^KO^* endothelium, correlating with disrupted cortical actin organization and dynamics.

### Scrib-actin clusters are present at the base of intact nascent sprouts and correlate with linear adherens junctions upon angiogenic stimulation

Scrib regulates integrity of tip cell-to-parent vessel adhesion (Fig. 2). We therefore next asked if Scrib-actin clusters function during angiogenic sprouting. We first investigated whether clusters are present at the sprout-parent vessel interface in Scramble and *SCRIB^KO^* microvessels. Upon angiogenic induction, we found that most Scramble microvessels exhibited at least 1 actin cluster at the sprout-parent vessel interface (Fig. 5f, g, arrows in inset). *SCRIB^KO^* microvessels occasionally exhibit actin enrichment at this interface (Fig. 5f, arrows in inset), but only rarely exhibit an actin cluster (Fig. 5f). Correlated with this decrease in actin clusters at the sprout-parent vessel interface, we also observed significantly decreased width of sprout bases from *SCRIB^KO^* microvessels (Fig. 5h). We then asked whether the organization of Scrib/actin clusters contributes to junction linearity in response to angiogenic stimulation. We applied vascular endothelial growth factor (VEGF) for 1 hour to Scramble and *SCRIB^KO^* monolayers cultured on compliant hydrogels. As previously reported, VEGF treatment caused an increase in stress fibers through the cell interior in both conditions (Fig. 5i) (Cao et al., 2017). In response to VEGF, *SCRIB^KO^* hMVECs exhibit markedly undulated junctions and lack cortical actin clusters, while in Scramble hMVECs, actin clusters remain present along with associated linear junctions (Fig. 5i). We conclude that Scrib-actin clusters are present at the base of most intact sprouts in Scramble microvessels, correlate with increased width of the base of nascent sprouts, and correlate with linear junctions in response to angiogenic chemokine.

### Scrib-enriched actomyosin clusters are contractile and regulate endothelial mechanics

Scrib organizes cortical actin clusters in endothelia, which are increasingly recognized as meso-scale organizers of the actomyosin cortex in other cell types (Akamatsu et al., 2014; Chou et al., 2024; Kruse et al., 2024; Luo et al., 2013). However, the composition of actin clusters and their functions coordinating endothelial cell mechanics remains poorly understood. To begin to understand how Scrib may organize clusters with consequences for broader endothelial cytoskeletal and adhesive organization, we examined potential components of clusters through immunofluorescence co-staining and Scrib co-immunoprecipitation. Clusters of Scrib and actin strongly overlap clustered structures of NMIIB, NMIIA, and phosphorylated myosin light chain (ppMLC, phosphorylated at Thr18/Ser19) (Fig. 6a). Notably, Scrib-actin clusters are distinct structures from the focal adhesion protein paxillin (Fig. S4e, f) and activated β1 integrin (CD29, Fig. S4g). Like actin, dynamic clustering events of NMIIB are observed in Scramble (Fig. 6b; Video 5); however, NMIIB clustering in *SCRIB^KO^* is severely disrupted (Fig. 6b; Video 5). Consistent with immunofluorescence overlap, we find that Scrib co-immunoprecipitates with NMIIB (Fig. 6c). Altogether, we conclude that Scrib plays a key role in organizing actin clusters, which contain contractile NMII but not focal adhesion components.

**Figure 6:**
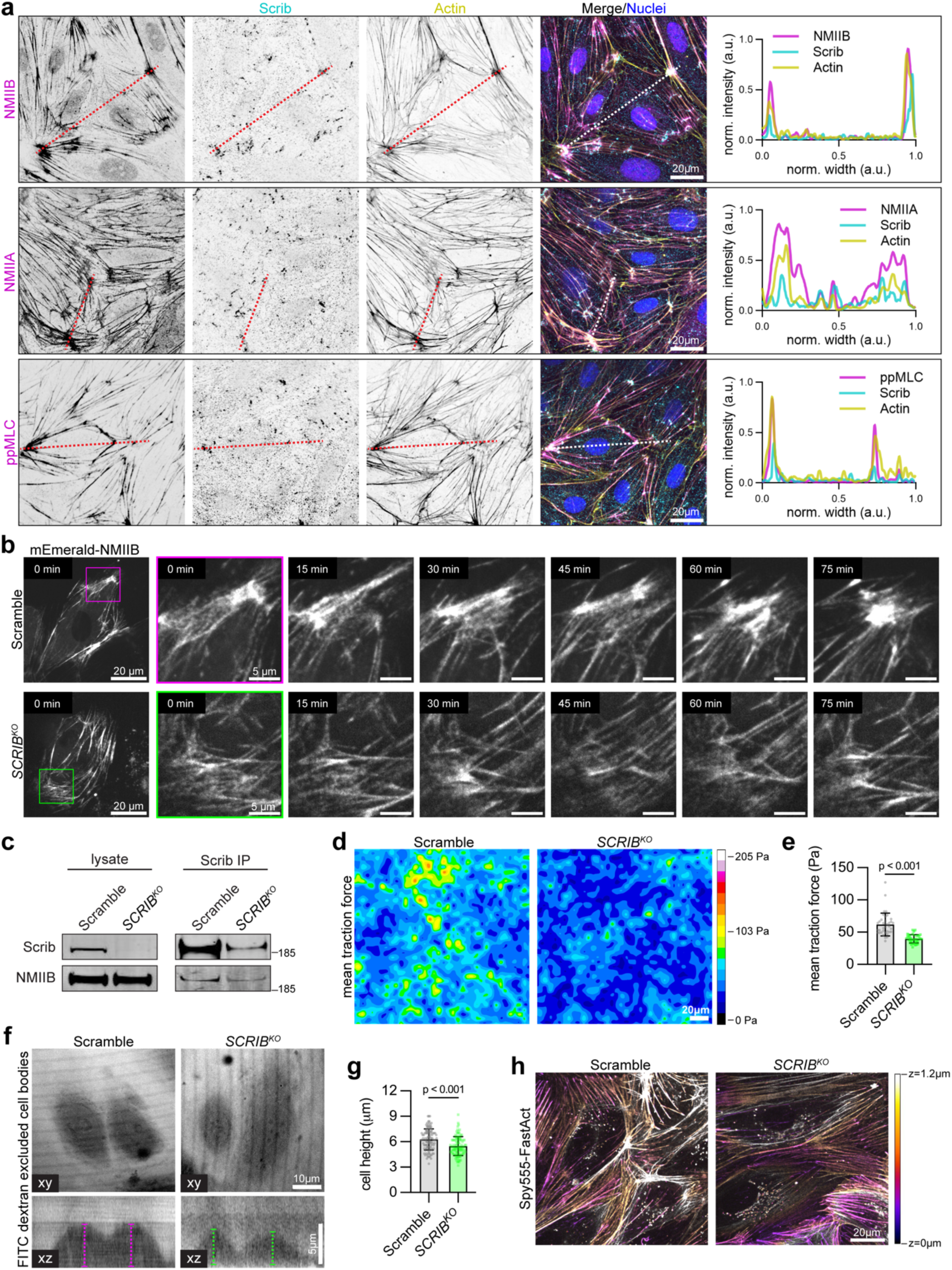
Scrib-enriched actomyosin clusters are contractile and regulate endothelial mechanics. **(a)** Representative micrographs of Scrib co-stained with F-actin, NMIIB, NMIIA, and pMLC at cortical clusters in control hMVECs, represented by line scans for each (N = 3 independent experiments). **(b)** Live time course of confluent monolayers heterogeneously expressing NMIIB (mEmerald-NMIIB) in Scramble and *SCRIB^KO^*. Inset of Scramble highlights formation of a large cluster over 75 min of retrograde NMIIB flow from the cell-cell adhesion. Inset of *SCRIB^KO^* highlights retrograde NMIIB flow from the cell-cell adhesion that fails to coalesce into a cluster (Video 5) (N = 3 independent experiments). **(c)** Immunoprecipitation of Scrib in Scramble and *SCRIB^KO^*, immunoblotted for Scrib and NMIIB (N = 3). **(d, e)** Traction force microscopy of confluent hMVEC monolayers cultured on compliant polyacrylamide gels. **(d)** Average intensity projection of force map and **(e)** quantification of mean traction force (N = 4 independent experiments, n ≥ 6 patches of 187 μm^2^ per condition per N, Mann-Whitney test). **(f)** Cell height measured through exclusion of FITC-dextran (70 kDa) in live cells, quantified in **(g)** (N = 3 independent experiments, n ≥ at least 12 cells per condition per N, 2-tailed student’s t-test). **(h)** Representative live actin images color-coded by z-position to highlight AJ-proximal cortical actin in Scramble and *SCRIB^KO^*. Abbreviations: Pa = Pascal.

Actomyosin clusters are important for integrating mechanical forces during wound closure and apical constriction (Bhat et al., 2024; Martin et al., 2009). We next sought to understand the function of Scrib-actomyosin clusters in regulating endothelial cell mechanics. We first measured traction across the cell-ECM interface, a measure of contractility in the actomyosin cortex (Oakes et al., 2012), and observed that *SCRIB^KO^* monolayers exert significantly less traction than Scramble monolayers (Fig. 6d, e). We also measured cell height as an overall read-out of polarity and actomyosin architecture in the cell (Pesen and Hoh, 2005; Rozman et al., 2020; Thiagarajan et al., 2022), using displacement of fluorescent 70 kDa dextran and confocal microscopy of confluent monolayers (Fig. 6f, g). Scramble hMVECs are on average 6.4 μm tall, while *SCRIB^KO^*hMVECs are significantly shorter at 5.5 μm (Fig. 6g). Most of the endothelial height is occupied by the nucleus, while the flat portion of the cell body where AJs and associated cortical actomyosin reside is only ∼1-1.5 μm in hMVECs (Fig. 6h). Using a live actin probe, confocal microscopy, and coding Z-depth by color, we find that the flat actin cortex is also shorter in *SCRIB^KO^* than Scramble. Interestingly, clusters in Scramble hMVECs tend to reside apically in Z-planes compared to other actin structures (Fig. 6h). Altogether, we find that loss of Scrib-actomyosin clusters is associated with broad changes in endothelial cortical mechanics, influencing cell-substrate traction forces as well as cellular architecture.

### *MYO1C^KO^* causes increased tip cell detachments and adherens junction undulation during angiogenic stimulation

Scrib organizes actomyosin clusters (Fig. 5, 6), in addition to regulating AJs (Fig. 3) and cortical actomyosin abundance (Fig. 4, 5) in endothelial cells, and these roles do not involve catenin-dependent cadherin coupling to actin (Fig. 4). We therefore asked whether another protein modulates the linkage of Scrib-actomyosin clusters to VE-cadherin. To test this hypothesis, we unbiasedly screened for interactors of Scrib and cross-checked these interactors with those identified by VE-cadherin-BioID (Fig. 1). We performed immunoprecipitation of Scrib and band excision mass spectrometry to identify proteins co-immunoprecipitating with Scrib (Fig. S5a). The most abundant protein identified was Myo1c, which is also highly abundant in the VE-cadherin-BioID (Fig. 1b). Myo1c is a molecular motor known to regulate epithelial AJs (Kannan and Tang, 2018; Tokuo and Coluccio, 2013) and insert into membranes with its tail domain and bind actin with its motor domain (McIntosh and Ostap, 2016). We validated the mass spectrometry result via immunoprecipitation of Scrib and immunoblotting for Myo1c (Fig 7a). We similarly confirmed the VE-cadherin-Myo1c interaction via immunoprecipitation of VE-cadherin and immunoblotting for Myo1c (Fig. 7b), which notably is maintained in *SCRIB^KO^*. Myo1c overlaps enriched actin structures and Scrib clusters (arrows) via immunofluorescence (Fig. 7c), while this localization is diminished in *SCRIB^KO^* (Fig. 7d, arrows). Importantly, junctional localization of Myo1c is maintained in *SCRIB^KO^*hMVECs (Fig. 7d, arrowheads).

**Figure 7:**
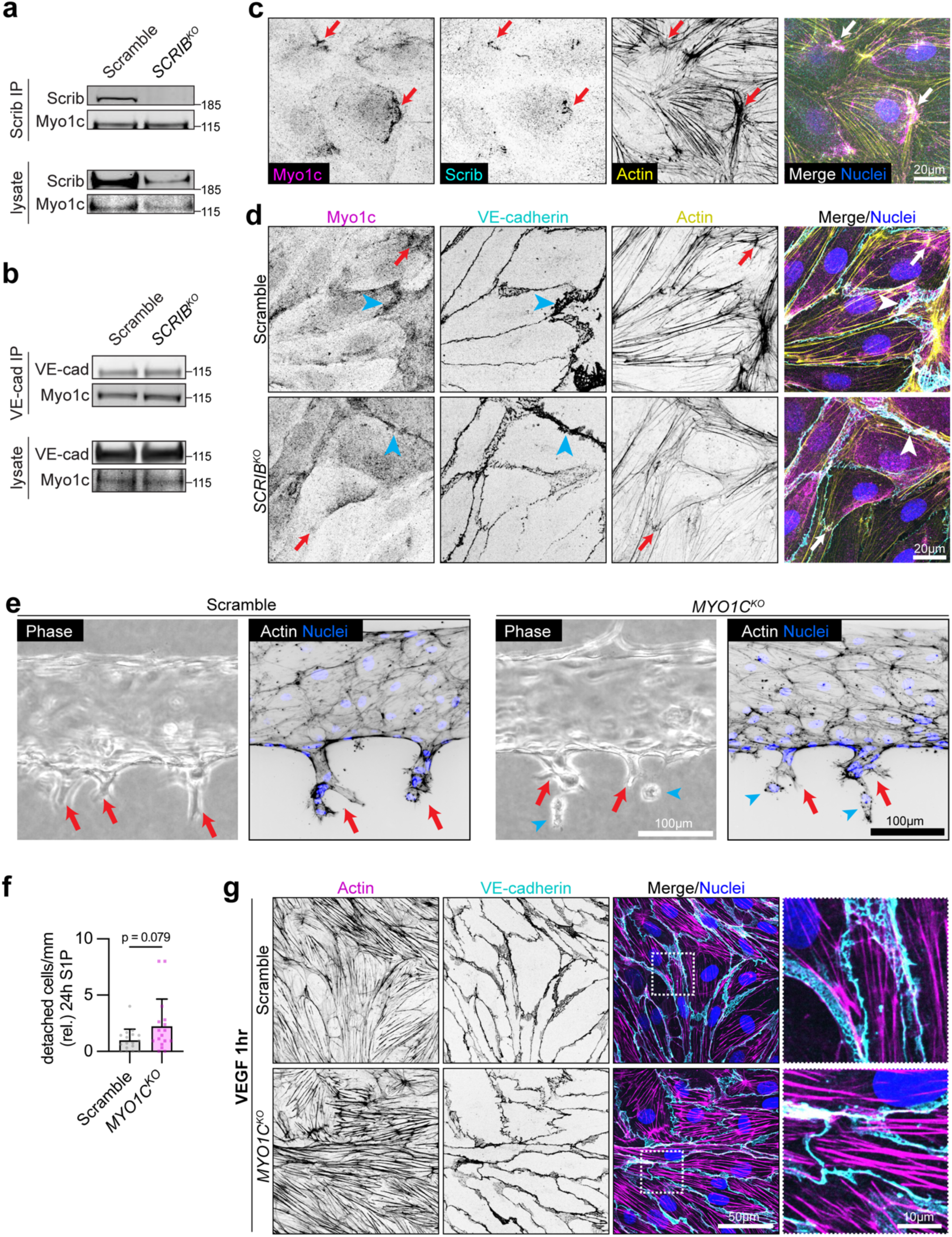
*MYO1C^KO^* causes increased tip cell detachments and adherens junction undulation with VEGF treatment. **(a)** Scrib immunoprecipitation in Scramble and *SCRIB^KO^*, immunoblotted for Scrib and Myo1c (N = 2). **(b)** VE-cadherin immunoprecipitation in Scramble and *SCRIB^KO^*, immunoblotted for VE-cadherin and Myo1c (N = 3). **(c)** Representative Myo1c immunofluorescence staining co-stained with actin and Scrib. Arrows denote areas of Myo1c intensity at Scrib/actin clusters (N = 2 independent experiments). **(d)** Representative Myo1c immunofluorescence staining in Scramble and *SCRIB^KO^* co-stained with actin and VE-cadherin. Blue arrowheads point to Myo1c overlap with VE-cadherin; red arrows point to Myo1c overlap with regions of enriched actin (N = 4 independent experiments). **(e)** Scramble and *MYO1C^KO^* microvessels stimulated with 500 μM S1P for 24 h, visualized by phase and fluorescence microscopy. Blue arrowheads indicate detached cells; red arrows indicate intact sprouts. Note: fluorescence images are not intensity matched to account for inter-vessel variability in actin staining. **(f)** Quantification of cell detachments in Scramble and *MYO1C^KO^*microvessels (N = 4 independent experiments, n ≥ 3 devices per condition per N, Mann-Whitney test). **(g)** Representative images of confluent Scramble and *MYO1C^KO^* endothelial monolayers grown on compliant polyacrylamide gels, treated with VEGF (50 ng/ml, 1 hr), and stained for actin and VE-cadherin (N = 3 independent experiments).

We next asked if Myo1c similarly plays a role in maintaining tip cell-to-parent vessel adhesion. We incorporated *MYO1C^KO^* hMVECs (Fig. S5b) into the same engineered model of angiogenesis in Fig. 2. Similar to *SCRIB^KO^*, we find that *MYO1C^KO^* form intact microvessels before S1P treatment (Fig. S5c, d) and exhibit a higher number of detached tip cells from the parent vessel than Scramble control vessels after S1P treatment (Fig. 7e, f; S5c, d). However, *MYO1C^KO^* does not disrupt actin clusters or cortex in homeostatic endothelium (Fig. S5e) like *SCRIB^KO^*. Since Myo1c association with actin and epithelial junctions increases with force (Greenberg et al., 2012; Kannan and Tang, 2018), we asked whether Myo1c might modulate the connection between Scrib-actomyosin clusters and VE-cadherin under angiogenic stimulation, when AJs are subject to increased forces (Barbacena et al., 2022; Yoon et al., 2019). When stimulated by VEGF, *MYO1C^KO^* cells exhibit highly undulated AJs (Fig. 7g), similar to *SCRIB^KO^* cells (Fig. 5i). Additionally, VEGF-stimulated *MYO1C^KO^* cells exhibit enriched actin stress fibers (Fig. 7g) and a small but significant decrease in Scrib cluster size compared to VEGF-stimulated Scramble cells (Fig. S5f, g). Together, we conclude that a Scrib-Myo1c axis links Scrib organization of the actomyosin cortex to VE-cadherin and maintains tip cell-to-parent vessel adhesion integrity and AJ linearity upon angiogenic stimulation.

## DISCUSSION

In this work, we demonstrate that Scrib regulates multicellular integrity during vascular sprouting at the early stage of tip cell-to-parent vessel adhesion maintenance. During angiogenic initiation, AJs loosen through a combination of actomyosin dynamics and internalization of VE-cadherin (Abraham et al., 2009; Cao et al., 2017; Grimsley-Myers et al., 2020) (Fig. 8b), and once a tip cell begins to extend (Fig. 8c), tip-stalk AJs experience an additional mechanical challenge (Barbacena et al., 2022; Yoon et al., 2019). Our data inform a model in which the junctional actomyosin cortex, reinforced and organized by a newly described mechanism involving Scrib-actomyosin clusters, is tethered in part via Myo1c to VE-cadherin. The AJs are sufficiently mechanically reinforced, and the tip cell-to-parent vessel connection remains intact (Fig. 8d). In contrast, during *SCRIB^KO^*tip cell emergence (Fig. 8c*), actomyosin clusters and junctional actomyosin are severely depleted, leading to decreased mechanical reinforcement of the AJ and ultimately tip cell detachment (Fig. 8d*). During *MYO1C^KO^*tip cell emergence (Fig. 8c**), there is decreased tethering between Scrib-actomyosin clusters and AJs leading to decreased reinforcement of the AJ and tip cell detachment (Fig. 8d**).

**Figure 8:**
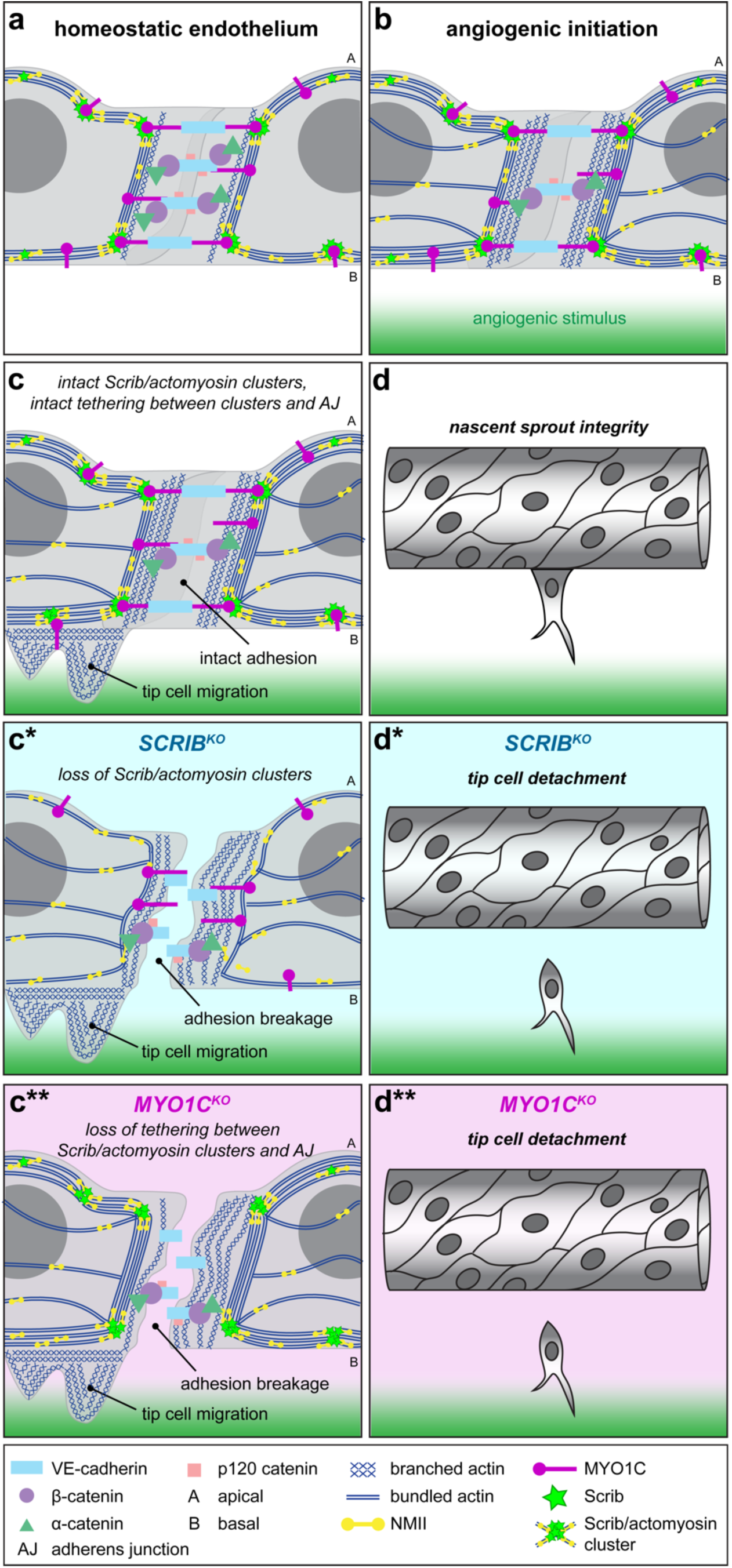
Summary model. **(a)** Scrib organizes cortical actomyosin clusters, which are critical for actomyosin cortex stability, including near AJs. Myo1c tethers these Scrib/actomyosin clusters to VE-cadherin and/or other cell membrane domains. **(b)** In a control endothelium exposed to angiogenic factors, junctions loosen through a combination of actomyosin dynamics and internalization of VE-cadherin. Cortical Scrib/actomyosin clusters mechanically stabilize cell-cell adhesions through interaction with Myo1c. **(c)** Once a tip cell is selected and begins to elongate and migrate towards the chemokine gradient, tip-stalk AJs experience an additional mechanical challenge. With Myo1c intact to tether Scrib/actomyosin clusters to VE-cadherin, junctions remain sufficiently mechanically reinforced, and the tip cell-to-parent vessel connection remains intact **(d)**. **(c*)** In a *SCRIB^KO^* endothelium undergoing angiogenic initiation, actomyosin clusters are severely depleted, and actomyosin broadly is less abundant in the junctional cortex. Myo1c can still localize to junctions and membrane, but the actomyosin cortex it tethers is not reinforced by actomyosin clusters. There is less mechanical reinforcement of the AJ by actomyosin clusters and cortex, which causes cell-cell adhesion destabilization, breakage, and a resulting tip cell detachment **(d*)**. **(c**)** In a *MYO1C^KO^* endothelium undergoing angiogenic initiation, there is a decrease in tethering between Scrib/actomyosin clusters and AJs. Similar to *SCRIB^KO^*, this loss of mechanical reinforcement of the AJ leads to junction destabilization, breakage, and tip cell detachment **(d**)**.

Scrib has critical roles in diverse cell types during tissue morphogenesis across organisms, including *Drosophila* epidermal cuticle formation (Bilder and Perrimon, 2000), vertebrate neural tube closure (Lesko et al., 2021; Murdoch et al., 2003; Robinson et al., 2012; Žigman et al., 2011), and mammalian ventricular myocardium septation (Boczonadi et al., 2014). Using an engineered angiogenic system that permits visualization of the earliest stages of angiogenesis, we find that Scrib regulates angiogenic sprout integrity, specifically at the stage of tip cell emergence from a parent vessel. This phenotype is reminiscent of recent work describing tip cell breakages following myosin II inhibition (Yoon et al., 2019), accompanied by AJ disorganization between tip and stalk cells. Similar tip cell breakages were observed upon deletion of Raf-1, which maintains Rho-kinase activity at tip cell AJs (Wimmer et al., 2012). Congruent with these observations, we observe that *SCRIB^KO^* causes VE-cadherin undulation and mislocalization of cortical actomyosin components. Junction remodeling is certainly required for angiogenesis (Abraham et al., 2009; Cao et al., 2017; Grimsley-Myers et al., 2020), but junction stability must be precisely tuned, even heterogeneously among neighboring cells (Bentley et al., 2014), to generate proper angiogenic sprouts. We posit that the mechanisms identified here contribute to a developing model in which Scrib essentially reinforces AJs via control of cortical actomyosin to ensure mechanical integrity of tip cell junctions.

We were surprised that Scrib stabilization of endothelial AJs was not explained by several Scrib mechanisms previously reported for epithelial AJs. Elegant work in Madin–Darby canine kidney (MDCK) cells shows Scrib depletion disrupts cell clustering via E-cadherin destabilization; this phenotype is partially rescued by expression of an E-cadherin-⍺-catenin fusion protein, leading to the conclusion that Scrib regulates the coupling between cadherin and catenins (Qin et al., 2005). In contrast, in endothelium, we did not observe a rescue of *SCRIB^KO^* cortical actomyosin or junction linearity by expression of a VE-cadherin-⍺-catenin fusion protein. A separate study in MDCK reported that Scrib regulates the interaction between E-cadherin and p120 catenin, and thereby E-cadherin endocytosis (Lohia et al., 2012). In endothelium, we observed that the interaction between VE-cadherin and p120 catenin is maintained in *SCRIB^KO^*, and *SCRIB^KO^* does not exhibit increased VE-cadherin endocytosis. A recent study in A549 lung carcinoma cells concluded Scrib depletion causes decreased ⍺-catenin and β-catenin localization at AJs (Abedrabbo et al., 2023). In *SCRIB^KO^* hMVECs, we observed no defects in ⍺/β-catenin localization at AJs nor a defect in β-catenin co-immunoprecipitation with VE-cadherin. Altogether, we provide evidence of a new mechanism in which Scrib regulates endothelial AJs and associated actomyosin separate from catenin-dependent cadherin coupling to actin, p120 catenin regulation of cadherin internalization, or interactions between VE-cadherin and catenins. These contrasting observations in epithelium suggest the possibility of alternative mechanisms for Scrib that converge on AJ stability.

Here we delineate a novel role for Scrib in organizing actomyosin clusters, which establish the architecture of the actomyosin cortex in endothelium. Previous work has described local regulation of NMII contractility by Scrib: Scrib scaffolds contractility regulators SGEF (Awadia et al., 2019) and DLC3 (Hendrick et al., 2016) at epithelial AJs. At the apical surface of intestinal epithelium, Scrib depletion disrupts phosphorylation of MLC (Boëda et al., 2023), and at the leading edge of migratory epithelium, Scrib depletion disrupts localization of Rho GTPases and p21-associated kinase (Dow et al., 2007; Nola et al., 2008) and atypical protein kinase C-associated NMIIB (Abedrabbo and Ravid, 2020). Our study uniquely demonstrates that *SCRIB^KO^*causes global disruption of myosin heavy chain filament architecture and dynamics due to loss of Scrib-organized clusters throughout the cortex. This suggests an architectural role for Scrib in assembling supramolecular NMII structures, rather than solely scaffolding effectors that regulate MLC phosphorylation. We find that actomyosin clusters stabilize endothelial AJs, but also maintain mechanics of the endothelial cortex broadly, including cortex height and traction forces, which may explain prior results implicating disrupted focal adhesions in *SCRIB^KO^* endothelium (Michaelis et al., 2013). Actomyosin clusters drive cell shape change in several morphogenic contexts, including *Drosophila* apical constriction (Martin et al., 2009), and mammalian epithelial wound closure (Bhat et al., 2024), though the composition and functions of actomyosin clusters are still being defined. Reconstituted NMII, actin, and actin cross-linkers can self-organize into nodal structures in vitro (Alvarado et al., 2013), but accessory proteins may modulate the size and dynamics of these structures (Kruse et al., 2024). Scrib may localize and organize clusters through multivalent interactions with NMII, as Scrib has multivalent protein-protein interaction domains and complexes with NMII here and in other studies (Abedrabbo et al., 2023; Abedrabbo and Ravid, 2020).

Whether this new mechanism for Scrib is cell-type specific, such as in the case of neurons where Scrib scaffolds proteins for synaptic vesicle clustering (Rui et al., 2017), or is conserved in other contexts, is unclear. One interesting possibility is that Scrib regulation of cortical actomyosin clusters is in fact conserved, and here is uniquely illuminated by the flat cell architecture of the endothelium. Several *SCRIB^KO^* phenotypes are conserved between epithelium and endothelium, such as such as decreased cell height (Boëda et al., 2023). The decreased cortical density of endothelium permits facile observation of actomyosin organization proximal to AJs (Efimova and Svitkina, 2018). Large actomyosin clusters are easily visible in endothelium, whereas some studies in mammalian epithelium have required treatment with latrunculin or blebbistatin for visualization (Luo et al., 2013). Interestingly, actomyosin clusters are associated with apical constriction, cell intercalation, and neural tube closure (Martin et al., 2009; Munjal et al., 2015; Nikolopoulou et al., 2019), morphogenic events that also require Scrib (Boëda et al., 2023; Lesko et al., 2021; Murdoch et al., 2003; Robinson et al., 2012; Žigman et al., 2011). Similar to our observations here, Scrib-depleted neural tube epithelium displays global disruption of NMII localization (Lesko et al., 2021). Additionally, Scrib is an established tumor suppressor in human cancers (Pearson et al., 2011; Zhan et al., 2008), and there is increasing recognition that disrupted cortical actomyosin tension contributes to epithelial tumorigenesis (Conti et al., 2024; Schipper et al., 2019). It is therefore plausible that direct maintenance of the actomyosin cortex could contribute to Scrib activities in the aforementioned settings.

Our work describes a novel mechanism by which Scrib, Myo1c, and VE-cadherin are coupled to ensure angiogenic sprout integrity. As Myo1c tethers actin to membranes (McIntosh and Ostap, 2016), we posit that Myo1c is a linker between cluster-reinforced actomyosin cortex and the plasma membrane, including at AJs. Unconventional myosin isoforms have been characterized within actomyosin clusters in yeast (Moseley and Goode, 2006). Recent work reported Myo1c and Scrib immunostaining in the same basolateral compartment of A431 epithelial cells at actin-rich protrusions (Troyanovsky et al., 2025). In our system, we observe Myo1c localization to clusters, though cluster organization is not largely dependent on Myo1c. We find that *MYO1C^KO^* causes junction undulation during angiogenic stimulation, and ultimately tip cell detachments from the parent vessel like *SCRIB^KO^*. This suggests that Myo1c maintains this crucial adhesion between tip cell and parent vessel via a VE-cadherin-Myo1c-Scrib complex. This is reminiscent to the role of Myo1c in epithelium, where it maintains E-cadherin-based AJs (Kannan and Tang, 2018; Tokuo and Coluccio, 2013), possibly through coupling between E-cadherin and actin (Kannan and Tang, 2018). In endothelium, Myo1c mediates force-intensive processes at the cell membrane, including exocytosis of large VWF cargo (El-Mansi et al., 2024). Myo1c is mechanosensitive and exhibits greater attachment to actin under tension (Greenberg et al., 2012) and increased localization to AJs under tension (Kannan and Tang, 2018). These functions are consistent with a potential role for Myo1c in stabilizing endothelial AJs during the mechanical perturbation caused by tip cell emergence.

Altogether, we reveal a novel role for Scrib in regulating endothelial mechanics and AJs via cortical actomyosin clusters, with the emergent role of maintaining nascent sprout integrity. Myo1c works in concert with both Scrib and VE-cadherin to facilitate this role, which is especially important during the mechanical perturbation on tip cell AJs during angiogenic initiation. This work underscores how actomyosin regulation of AJ dynamics is necessary for initial angiogenic sprouting but is precisely controlled to allow proper multicellular vessel growth. A deeper understanding of this process may present an opportunity for better therapeutic control of vascularization.

## Supporting information

Supplemental Table 1: CRISPR Primers

Supplemental Table 2: Primers for DNA Fragment Assembly

Supplemental Table 3: VE-cadherin BioID

Supplemental Table 4: Abbreviations

Source Data 1: Full Blots

Source Data 2: Source Data for Graphs

## Acknowledgements

This work was supported by National Institutes of Health (NIH) grants R35GM150987 (M.L. Kutys), F31HL172606 (L.N. Mayo), shared equipment grant S10OD028611 (Nikon CSU-W1 SoRa), and UCSF/UCB Joint Graduate Group in Bioengineering T32GM139794 (F. Duong). Other support was from the Leducq Foundation 21CVD03 (M.L. Kutys and M.L. Iruela-Arispe), Tobacco-Related Disease Research Program Predoctoral Fellowship T33DT6442 (K.A. Jacobs) and National Science Foundation Graduate Research Fellowship 2038436 (K.A. Jacobs). The authors thank Jon Humphries for assistance with VE-cadherin-BioID. We also thank the following for their critical insights on this project: Tania Singh, Diane Barber, Torsten Wittmann, Jeffrey Bush, Orion Weiner, Xiaohua Gong, Dean Sheppard, and Rong Wang.

## METHODS

### Cell culture

Primary human dermal blood microvascular endothelial cells (hMVECs) were obtained from human neonatal foreskin (Lonza). hMVECs were maintained from passages 2-7 in EGM2-MV medium (Lonza) and passaged at 80-90% confluency. Unless otherwise mentioned, end-point experiments (e.g. western blot lysis or fixation for immunofluorescence) were cultured until cells reached 100% confluency. HEK-293T cells (Clonetech) were maintained from passages 3-20 in high glucose DMEM (Sigma) supplemented with 10% fetal bovine serum (Peak or SeraPrime), 2 mM L-glutamine (Invitrogen), 1 mM sodium pyruvate (Gibco) and 1% penicillin/streptomycin (Sigma). All cells were maintained in a humidified incubator (Thermo Fisher Scientific) at 37C and 5% CO_2_ and tested regularly for mycoplasma (Applied Biological Materials). During passaging, cells were counted using a Trypan Blue stain (Invitrogen) and the Countess 3 automated cell counter (Invitrogen).

### Antibodies and reagents

Antibodies against GAPDH (14C10, 1:10,000 WB), pMLC Ser19 (3675, 1:300 IF, 1:1,000 WB), ppMLC Thr18/Ser19 (95777S, 1:300 IF, 1:1,000 WB), and p120 catenin (4989, 1:300 IF, 1:1,000 WB) were from Cell Signaling Technologies. Antibodies against CD29 (activated β1 integrin) (9EG7, 1:300 IF) and β-catenin (14, 1:300 IF, 1:1,000 WB) were from BD Biosciences. Antibodies against Myo1c (HPA001768, 1:300 IF, 1:1,000 WB) and Scrib (HPA023557, 1:300 IF, 1:1,000 WB) were from Human Protein Atlas. Antibodies against non-muscle myosin IIA (19098, 1:300 IF, 1:5,000 WB) and non-muscle myosin IIB (19099, 1:300 IF, 1:5,000 WB) were from BioLegend. Antibodies against Paxillin (610620, 1:300 IF) and ⍺-catenin (AB_397593, 1:1,000 WB) were from BD Transduction Laboratories. Antibodies against Scrib (C-6, 1:300 IF, 1:1,000 WB) and VE-cadherin (F-8, 1:200 IF, 1:1,000 WB) were from Santa Cruz Biotechnology. Antibody against VE-cadherin (ab33168, 1:300 IF, 1:1,000 WB) was from Abcam. Antibody against ⍺-catenin (C2081, 1:300 IF, 1:1,000 WB) was from Sigma Aldrich. Non-blocking antibody against VE-cadherin ECD conjugated to Alexa Fluor 647 (55-7H1, 1:200 live, 1:200 IF) was from ThermoFisher. Rhodamine phalloidin (1:2,000), Alexa Fluor 488 conjugated phalloidin (1:400), and Alexa Fluor 488, 568, and 647 goat anti-mouse and anti-rabbit IgG secondary antibodies (1:400) were from Invitrogen. IRDye donkey 680 and 800 antibodies (1:10,000 WB) were from LICOR Bio. DAPI nuclear stain (1:400) and FITC Dextran 70kDa (1:1,000 live) were from Sigma. Sphingosine-1-phosphate was from Tocris Bioscience (500nM). Human vascular endothelial growth factor was from Peprotech (50ng/ml). Spy555-FastAct (1:2,000) was from Cytoskeleton Inc.

### Lentiviral-mediated CRISPR-Cas9 editing

Primary human microvascular cells (hMVECs) were edited via lentiviral-mediated CRISPR-Cas9 editing as described previously (Kutys et al., 2020; Polacheck et al., 2017). Briefly, single guide RNAs (sgRNAs) for the protein of interest were designed using CRISPOR (Concordet and Haeussler, 2018) or E-CRISP (Heigwer et al., 2014). Primers for these sgRNAs (top and bottom strands, Supplementary Table 1) were annealed with T4 polynucleotide kinase (T4 PNK, NEB) then ligated into the BsmBI site of pLentiCRISPRv2 (plasmid #52961; Addgene) with T4 ligase (NEB). This plasmid was transfected via calcium phosphate transfection into HEK-293T cells alongside packaging plasmids psPAX2 (plasmid #12260; Addgene) and pMD2.G (plasmid #12259; Addgene). After 48 h, viral media was harvested and filtered through a 0.22 μm sterile syringe filter. Lentivirus was concentrated via incubation with 1x polyethylene glycol lentiviral concentrator solution (3x solution comprised of 10% w/v Polyethylene Glycol 8000 and 0.3 M NaCl) for at least 4 h on a rotator at 4C, then centrifugation at 4C and 1600g for 1 h. The viral pellet was resuspended in sterile PBS, then stored as aliquots at -80C. ∼150,000 hMVECs were transduced with 50-100 μl of lentivirus for 12-18 h. After 3 days in culture, hMVECs were selected using 2 μg/mL puromycin for 4 days, then lysed for Western blot to confirm the knockout. Once confirmed, CRISPR KO primary cell stocks were expanded and froze down at p4 or p5 to use for experiments.

### Lentiviral-mediated gene expression

All fusion proteins were assembled into a lentiviral pRRL vector at the KpnI and EcoRI sites and expressed using a CMV promoter. Primers to generate these constructs were designed using NEBuilder and SnapGene Viewer and are listed in Supplementary Table 2. DNA fragments were assembled using NEBuilder HiFi DNA Assembly Master Mix. mEmerald-MyosinIIB was a gift from M. Davidson’s lab. For the VE-cadherin-⍺-catenin-mEmerald chimera fusion protein, fragments of mouse VE-cadherin and ⍺-catenin were isolated from mouse cDNA library, and mEmerald was a gift from M. Davidson’s lab. For LifeAct-mScarlet, LifeAct was isolated from LifeAct-GFP (plasmid # 51010, Addgene), and mScarlet was a gift from Y. Liu. For VE-cadherin-BioID-HA and BioID-HA, human VE-cadherin was a gift from M. Davidson’s lab, and BioID-HA was isolated from pcDNA3.1 MCS-BirA(R118G)-HA (plasmid #36047). pRRL-based plasmids were tranfected via calcium phosphate transfection into HEK-293T cells alongside packaging plasmids psPAX2 (plasmid #12260; Addgene) and pMD2.G (plasmid #12259; Addgene). After 48 h, viral media was harvested and filtered through a 0.22 μm sterile syringe filter. Lentivirus was concentrated via incubation with PEG lentiviral concentrator solution (4x Polyethylene Glycol 8000, Fisher) for at least 4 h on a rotator at 4C, then centrifugation at 4C and 1600 g for 1 h. The viral pellet was resuspended in sterile PBS, then stored as aliquots at -80C. hMVECs were transduced for 12-18 h before media change and cultured for at least 2-3 days to allow robust protein expression before use in experiments.

### VE-cadherin BioID

VE-cadherin-BioID (VE-BirA*) and cytoplasmic-BioID (cyto-BirA*) lentiviral constructs were utilized as described previously (Polacheck et al., 2017). hMVECs stably expressing VE-BirA* or cyto-BirA* were generated via lentiviral transduction. For each condition, EGM2-MV containing 50 µM biotin was added to two confluent 15 cm plates for 20 hours. After incubation, hMVEC monolayers were lysed in RIPA buffer (pH 7.4, 50 mM Tris, 150 mM NaCl, 1% v/v Triton-X, 0.1% w/v sodium dodecyl sulfate, 0.5% w/v sodium deoxycholate) and biotinylated proteins were extracted by incubating lysates with Dynabeads MyOne™ Streptavidin C1 (Invitrogen) beads for 1 hour at room temperature. Beads were washed three times in RIPA buffer and biotinylated proteins were denatured and eluted in 2X sample buffer containing biotin at 95C for 10 minutes. Samples were isolated by gel top SDS-PAGE extraction. In-gel tryptic digestion was performed, and peptides were desalted using POROS Oligo R3 beads (Thermo Fisher Scientific). Peptides were washed with 0.1% formic acid and peptides were eluted with 50% acetonitrile and 0.1% formic acid. Peptides were dried via lyophilization and resuspended in 5% acetonitrile and 0.1% formic acid before analysis by liquid chromatography-tandem mass spectrometry (LC-MS/MS) by MS Bioworks. The complete list of identified proteins is provided in Supplementary Table 3. Proteins were sorted by abundance of spectra for VE-BirA* and highlighted if the number of spectra for Cyto-BirA* was less than 20.

### Immunoblotting

hMVECs were cultured to 100% confluency on a 6 well plate, 1 well per condition. Cells were washed with cold 1x PBS plus calcium and magnesium (0.9 mM CaCl_2_, 0.49 mM MgCl_2_), lysed in cold RIPA lysis buffer (pH 7.4, 50 mM Tris, 150 mM NaCl, 1% v/v Triton-X, 0.1% w/v sodium dodecyl sulfate, 0.5% w/v sodium deoxycholate) with 2x protease/phosphatase inhibitor (Thermo), scraped and pipetted into an Eppendorf tube, and incubated on ice for 20 min. Lysates were then centrifuged for 10 min at 4C at 21,380 g (SCILOGEX). Supernatants were isolated from the pellet, flash frozen with liquid N_2_, and stored at -80C if not used immediately. Protein concentration in the lysate for each condition was determined using a bicinchoninic acid (BCA) reaction kit (Prometheus). Lysate was denatured using 1x NuPAGE LDS Sample Buffer (Life Technologies) containing 5% β-mercaptoethanol (Sigma) for 10 min at 70C. Denatured lysate with equalized concentrations of protein and PageRuler prestained protein ladder (Thermo) were loaded into Invitrogen NuPAGE 4 to 12% BisTris protein gels and run at 140-160V in 1x MOPS/SDS running buffer (20x MOPS/SDS buffer comprising 1 M MOPS, 1 M Tris base, 69.3 mM sodium dodecyl sulfate, 20.5 mM EDTA free acid). Proteins were transferred from the gel onto polyvinylidene fluoride (PVDF) membrane (Millipore) or nitrocellulose membrane (NETA) in cold 1x transfer buffer with 10% methanol (20x transfer buffer comprising 50 0mM bicine, 500 mM Bis-Tris, 20.5 mM EDTA free acid) using a Mini Trans-Blot Cell (Bio-Rad) at 4C. Membranes were blocked in 5% nonfat milk (Apex) or 5% bovine serum albumin (Fisher) in TBST (1x TBS + 0.1% Tween). Primary antibody was suspended in blocking solution and incubated for 1-2 h at RT with gentle rocking. Membranes were washed 3x in TBST for 10 min each with gentle rocking. Secondary antibody was suspended in blocking solution and incubated for 1 h at RT with gentle rocking. Membranes were washed 3x in TBST for 10 min each with gentle rocking. Membranes were imaged on an Odyssey CLx LICOR Imaging System and quantified using either LICOR software or ImageJ. Immunoblots were adjusted for brightness and contrast using ImageJ, and intensity values were normalized to the GAPDH loading control and biological replicate control (e.g. Scramble). Full, uncropped Western blots are provided in Source Data files.

### Immunoprecipitation and mass spectrometry

hMVECs were cultured to 100% confluency on a 10 cm plate, 1 plate per condition. Cells were washed with cold 1x PBS plus calcium and magnesium (0.9 mM CaCl_2_, 0.49 mM MgCl_2_), lysed in cold IP lysis buffer (pH 7.4, 25 mM Tris, 150 mM NaCl, 5 mM MgCl_2_, 1% v/v Triton-X) with 2x protease/phosphatase inhibitor (Thermo), scraped and pipetted into an Eppendorf tube, then needle lysed 20x using a 1 ml syringe and 22 gauge needle. Lysates were incubated on ice for 10 min then centrifuged for 10 min at 4C at 21,380 g (SCILOGEX). Supernatants were isolated from the pellet, flash frozen with liquid N_2_, and stored at -80C if not used immediately. Protein concentration in the lysate for each condition was determined using a bicinchoninic acid (BCA) reaction kit (Prometheus). Concentration of protein was equalized across conditions for all steps. A small amount of lysate for each condition was conserved separately from the immunoprecipitation tube to run as input on the same gel. For immunoprecipitation, primary antibody was added to lysate tubes at a ratio of 2 or 3 μg antibody per 1 mg of protein in lysate. This solution was incubated on a tube rotator for 2 h at 4C. Then, the antibody/lysate solution was incubated with 50 μL Pierce Protein A/G beads (Thermo) on a tube rotator for 2 h at 4C. Beads were spun down and washed 3x with IP lysis buffer. Lysate tubes and bead tubes were denatured using 2x NuPAGE LDS Sample Buffer (Life Technologies) containing 5% β-mercaptoethanol (Sigma) for 10 min at 70C. For immunoprecipitation and immunoblot, immunoblot was conducted exactly as described in “Immunoblotting” section. For immunoprecipitation and Coomassie stain, the lysates were run on a gel as described in the “Immuoblotting” section but stained for 1 hour in a 15 cm plate with Novex Simply Blue Safe Stain (Thermo) and washed overnight with water. Bands were visualized on the Odyssey CLx LICOR Imaging System and excised with a clean razor blade if sending to mass spectrometry. Single, excised bands for protein identification were analyzed by LC-MS/MS by MS Bioworks as follows: In-gel digestion with trypsin was performed using a DigestPro robot (CEM). Gel bands were washed with 25 mM ammonium bicarbonate followed by acetonitrile, reduced with 10 mM dithiothreitol at 60C followed by alkylation with 50 mM iodoacetamide at RT, digested with trypsin (Promega) at 37C for 4 h, and quenched with formic acid, and the supernatant was analyzed directly without further processing. Half of the digested sample was analyzed by nano LC-MS/MS with a Waters M-Class HPLC system interfaced with a ThermoFisher Fusion Lumos mass spectrometer. Peptides were loaded on a trapping column and eluted over a 75 μm analytical column at 350 nL/min; both columns were packed with Luna C18 resin (Phenomenex). The mass spectrometer was operated in data-dependent mode, with the Orbitrap operating at 60,000 FWHM and 15,000 FWHM for MS and MS/MS, respectively.

### Microfluidic device design and fabrication

*Device design.* The angiogenic device design was adapted from previously published protocols (Nguyen et al., 2013; Wang et al., 2020). Detailed device designs with relevant dimensions are provided in Supplementary Files. Device master layers were designed in Adobe Illustrator. The device described uses 3 layers, each containing the ECM area and media ports. The base layer (layer 1) creates a needle buffer of 120 μm in height to prevent the acupuncture needle from touching the glass coverslip. The middle layer (layer 2) is designed to accommodate the acupuncture needle with a height of 160 μm. The top layer (layer 3) contains only the ECM area and media ports to add height to these features with a height of 120 μm.

*Photolithography.* Layers were fabricated using photolithography as previously described (Polacheck et al., 2019). A silicon wafer was surface functionalized with a plasma cleaner (Harris Plasma) at 0.3 Torr for 5 min. SU-8 2010 (Kayaku Advanced Materials) was deposited onto the surface of the wafer at desired thicknesses using a spin coater (Laurell) and baked pre- and post-UV exposure at 1.5x the recommended lengths of time detailed in the SU-8 protocol. Each layer was patterned via UV exposure using the Alveole system and a Nikon Ti2 microscope at 4x magnification, using recommended power settings in the SU-8 protocol. Between layer 1 and layer 2, a blocking layer was spun and baked onto the wafer to prevent UV exposure of layer 1’s needle buffer during exposure of layer 2. The blocking layer was composed of 70 mL of SU-8 2010 and 30 mL of S1813 (MicroChem), mixed overnight at 37C on an orbital shaker. For exposure of layer 2 and 3, epifluorescence light was used to align the pattern to layer 1’s pattern so that each layer was patterned exactly on top of the previous. After all layers were fabricated, unpolymerized SU-8 was removed via incubation with propylene glycol monomethyl ether acetate (PGMEA, Sigma) for at least 1.5 h with gentle agitation, replacing with fresh PGMEA as necessary. After drying the wafer with compressed air, the wafer was surface treated with trichloro(1H,1H,2H,2H-perfluorooctyl)silane (Sigma) via vapor deposition overnight to prevent polydimethylsiloxane (PDMS Sylgard 184, Dow Corning) from binding to the wafer.

*Soft lithography.* Soft lithography with PDMS was used to generate copies of the microfluidic device. First, the wafer was attached to the base of a 10 cm petri dish with double sided tape. 40 g PDMS base was mixed 10:1 by weight with PDMS curing agent (Sylgard 184, Dow Corning), degassed with vacuum desiccation, then poured on the wafer. This initial PDMS pour was polymerized overnight at 60C, then used as a “master” to create a sturdy plastic replica of the wafer (Smooth-Cast 305, Smooth-On), on which all further PDMS was poured to create device copies.

*Device assembly and surface treatment.* Device preparation was adapted from previously published protocols (Pérez-Rodríguez et al., 2021; Polacheck et al., 2019; Rathod et al., 2024). Devices were cut from the white plastic master using a razor blade. Biopsy punches (Thermo or Electron Microscopy Sciences) were used to cut the media ports (6 mm) and ECM ports (1.5 mm) from each device, then devices were cut into individual pieces using a razor blade. 24 mm x 24 mm glass coverslips (Sigma) and devices were cleaned with lab tape and plasma ashed for 30s at 0.3 Torr. Individual devices were pressed onto coverslips and permanently bound by baking at 100C for 10 min. A solution of 2 mg/ml dopamine hydrochloride (Sigma) was prepared in 10 mM Tris, pH 10 and pipetted into the ECM ports of the device. This surface treatment was incubated for 1 h protected from light, then washed with dH2O 3x before bathing the devices in H2O in a beaker for at least 1 h.

*Device sterilization and needle insertion.* In preparation for cell seeding within the device, the devices were bathed in 70% ethanol for at least 30 min. Steel acupuncture needles of 160 μm diameter (Tai Chi) were sterilized in 70% ethanol in a bench top ultrasonic cleaner (Sper Scientific), dried within a biosafety cabinet, then incubated in sterile filtered 0.1% bovine serum albumin (Fisher) for 30 min. Both devices and needles were allowed to dry in a biosafety cabinet, then needles were inserted into the two device channels. Devices were further sterilized by exposure to UV in a PCR chamber (Plas Labs) for 15 min. The assembled devices were dried completely in a vacuum desiccator for at least 1 h, up until immediately before addition of collagen I to the ECM chamber.

*Collagen addition and needle removal.* Rat tail collagen I (Corning) was brought to a final concentration between 2.5 and 3 mg/ml at pH 7.8 on ice. To accomplish this, 10x DMEM (Thermo) and 10x reconstitution buffer (1.2 g of NaHCO3 and 4.8 g of HEPES in 50 mL H2O) were mixed on ice with a titrated amount of 1 N NaOH, then collagen was added and mixed carefully to avoid introducing bubbles. Devices were removed from the desiccator, then collagen was pipetted into an ECM port of each device to fill the ECM chamber. The devices were immediately moved to the incubator for 30 min. After this, fresh media was added to the media ports of each device and a PBS-soaked sterile Kimwipe was added to the petri dishes to maintain humidity. After at least 4 h, needles were removed from the device using tweezers, and sterile vacuum grease (Dow Corning) was used to seal the needle entry points. Fresh media was added, and the devices were placed on a laboratory rocker at 5 rpm +/- 30° tilt for at least 8 h.

### Engineered 3D angiogenesis model culture

*Cell seeding.* For each device, one channel and associated media ports were designated as “cell channel” (typically the “top” channel as shown in Figure 2), while the other channel and associated media ports were designated as “blank channel” (bottom channel as shown in Figure 2). Endothelial cells were passaged and resuspended at 1x10^6^ cells/ml. For each device, media was aspirated gently from each media port. 40 μl media was added to each of the bottom media ports. 40 μl cell suspension was added to the top left media port, then 45 μl cell suspension was added to the top right media port to drive gentle fluid flow in the leftward direction. This flow permitted cells to move from the top right media port into the channel and adhere to the collagen walls. For roughly half the time, the petri dish was flipped so that cells would seed the top and bottom of the channel evenly. Once seeded at an appropriate density (Polacheck et al., 2019), cell suspension and media was removed from all wells and replaced with 100 μl fresh media. At least three devices were prepared for each condition tested.

*Device maintenance and culture.* Devices were continuously cultured on a laboratory rocker at 5 rpm +/- 30° tilt in a humidified incubator. Media was changed on the devices every 24 h (100 μl per media port). Typically 24-48 h of culture was required to obtain fully endothelialized channels.

*Preparation for endpoint experiments.* Once devices were fully endothelialized in the top channel, angiogenic factors could be introduced into the bottom channel. This typically occurred 36-48 h post seeding. To drive sprouting angiogenesis, 500 μM S1P was prepared in culture media and added to the bottom channel, while untreated culture media was added to the top/cell channel. For immunofluorescence and sprout parameter quantification, the “endpoint” was considered 24 h of treatment. Phase images were taken immediately prior to S1P treatment and immediately after 24 h of S1P treatment using an Olympus inverted microscope CKX53, Ritega R1 CCD camera, 10x phase objective, and Olympus EPView software. Devices were fixed in 4% paraformaldehyde (Electron Microscopy Sciences) prepared in 1x PBS plus calcium and magnesium (0.9 mM CaCl_2_, 0.49 mM MgCl_2_) pre-warmed to 37C. During fixation, devices were placed on the same laboratory rocker for culture at 37C for 20 min, then washed with 1x PBS. For live imaging of sprouting events, devices were cultured with S1P for 12 h on the laboratory rocker, then moved to a microscope (Nikon Ti2, 10x objective, Oko humidified live imaging chamber, Nikon Elements software) for live phase imaging for 12h. Further microscope details are available in “Live-cell microscopy”.

### Hydrogel coverslip preparation

Compliant polyacrylamide (PA) gels of approximate stiffness 2.5 kPa were synthesized on 18mm glass coverslips for use in immunofluorescence and traction force experiments. First, glass coverslips (18 mm or 25 mm diameter) were cleaned and activated via plasma ashing at 0.3 Torr. Coverslips were further activated by the following steps gently on an orbital shaker: incubating with 0.5% v/v (3-Aminopropyl)triethoxysilane (APTES, Thermo) in H_2_O for 30 min, washing 3x with H2O for 10 min each, incubating with 0.5% v/v glutaraldehyde (Thermo) for 1 h, washing 3x with H2O for 30 min, and allowing to dry between two Kimwipes overnight (Syga et al., 2018).

Then, polyacrylamide gels were polymerized on the activated coverslip surface. Quartz glass slides (Thermo) were passivated overnight via vapor deposition of trichloro(1H,1H,2H,2H- perfluorooctyl)silane (Sigma), then cleaned with distilled H_2_O and placed in a 150mm petri dish. The gel solution was mixed in a 1.7 μl tube: 125 μl 40% acrylamide, 100 μl 2% bis-acrylamide (Bio-Rad), 100 μl 10x PBS, 574 μl H_2_O, 1 μl TEMED (Bio-Rad), and 100 μl 1% v/v ammonium persulfate in H2O (Shebanova, 2010). For traction force microscopy, 200nm green fluorescent beads (Thermo) were mixed at 1:100 in H_2_O, and this H_2_O was used for the 574 μl component. Immediately after mixing, drops of gel solution were added to the quartz glass slides (20 μl for 18mm coverslips, 50 μl for 25mm coverslips). Coverslips were applied, activated side facing the drop, onto the drop using forceps. Gels polymerized in a dish humidified by PBS-soaked Kimwipes for 45-60 min. Distilled H_2_O was added to submerge the quartz-gel-coverslip sandwiches for at least 5 min, then gel-coverslip was gently removed from the passivated quartz using a fresh razor blade.

Next, the gel surface was activated using a solution of L-dopa (2 mg/ml, Thermo) in 10mM Tris, pH 8.5. L-dopa was allowed to dissolve for 45 min on a tube rotator protected from light, then sterile filtered. A square of fresh Parafilm was stuck to a 15 mm dish, and drops of L-dopa were added to the surface (50 μl for 18mm coverslip gels, 100 μl for 25mm coverslip gels). Coverslip gels were flipped onto these drops, gel side facing the drop. This incubated for 75 min protected from light, then coverslip gels were washed in distilled H_2_O for at least 5 min in a 15 cm petri dish. After washing with dH2O, coverslip gels were incubated with a solution of 50 μg/ml collagen I (Corning) in 1x PBS for 2 h at 37C. Coverslip gels were then washed with sterile 1X PBS and sterilized using a PCR UV chamber (Plas Labs) for 10 min. Coverslip gels were washed twice more with sterile PBS before use in cell culture.

### VE-cadherin internalization assay

VE-cadherin internalization assay was conducted according to previously published work (Malinova et al., 2021). Confluent hMVECs seeded onto compliant polyacrylamide gel coverslips were incubated with an Alexa Fluor-647-conjugated non-blocking antibody against the extracellular domain of VE-cadherin (55-7H1) diluted 1:200 in culture media for 30 min at 4C to block internalization. Then, cells were washed with 1x PBS twice and incubated in unlabeled culture media for 2 h. Fixation was conducted with 4% paraformaldehyde (Electron Microscopy Sciences) prepared in 1x PBS plus calcium and magnesium (0.9 mM CaCl_2_, 0.49 mM MgCl_2_) pre-warmed to 37C, incubated on coverslips for 10 min at 37C. Coverslips were washed with 1x PBS and mounted in 15 μl ProLong Glass Antifade Mountant (Invitrogen) on standard microscope cover glasses. Coverslips were sealed with clear nail polish (Electron Microscopy Science) and stored at 4C. VE-cadherin internalization was assessed by analyzing maximum intensity projections of VE-cadherin post-internalization assay in ImageJ. Cells were assigned a number for each field of view. Each cell was duplicated into a new image, outlined, and the junction/cell exterior were cleared. For each biological replicate, a constant threshold value was utilized to visualize discrete particles without including background staining and was used to create a mask of VE-cadherin+ vesicles within the cell. The watershed function was applied as necessary to separate clusters of vesicles. “Analyze Particles” was used to quantify particles between 0.2 and 10 μm^2^ with circularity 0.1-1. Number of particles per cell was quantified. For each biological replicate, values were normalized to the mean Scramble value of particles per cell.

### Immunofluorescence and confocal microscopy

#### 2D coverslip immunofluorescence staining

Coverslips were fixed when hMVECs reached 100% confluency. Fixation was conducted with 4% paraformaldehyde (Electron Microscopy Sciences) prepared in 1x PBS plus calcium and magnesium (0.9 mM CaCl_2_, 0.49 mM MgCl_2_) pre-warmed to 37C, and incubated on coverslips for 10 min at 37C. Coverslips were washed once with 1x PBS, then quenched with 100 mM glycine (Sigma) in H_2_O for 1 h at RT. Coverslips were permeabilized with 0.1% Triton-X (Sigma) in 1x PBS for 10 min. Coverslips were washed twice in PBS, then blocked in a solution of 5% bovine serum albumin (Fisher) and 4% normal goat serum (Millipore) in 1x PBS for 1 h at RT or overnight at 4C. Primary antibodies were prepared in blocking solution at concentrations detailed in the antibody section. Coverslips were incubated with primary antibodies for 1-2 h at RT, then washed 3 times with 1x PBS for 10 min each. Secondary antibodies and fluorescent dyes were prepared in blocking solution at concentrations detailed in the antibody section. Coverslips were incubated with the solution of secondary antibodies and fluorescent dyes for 1-2 h at RT protected from light, then washed 3 times with 1x PBS for 10 min each. Coverslips were mounted in 15 μl ProLong Glass Antifade Mountant (Invitrogen) on standard microscope cover glasses. Coverslips were sealed with clear nail polish (Electron Microscopy Science) and stored at 4C.

#### 3D microvessel immunofluorescence staining

Microvessel devices were fixed in 4% paraformaldehyde (Electron Microscopy Sciences) prepared in 1x PBS plus calcium and magnesium (0.9mM CaCl_2_, 0.49mM MgCl_2_) pre-warmed to 37C. During fixation, devices were placed on the same laboratory rocker for culture at 37C for 20 min. All remaining steps took place on a laboratory rocker with channels positioned parallel to the rocking axis at RT. Post fixation, devices were washed twice with 1x PBS for 10 min each, then quenched with 100mM glycine (Sigma) in H_2_O for 1 h at RT. Devices were permeabilized with 0.25% Triton-X (Sigma) in 1x PBS for 20 min. Devices were washed twice in PBS, then blocked in a solution of 5% bovine serum albumin (Fisher) and 4% normal goat serum (Millipore) in 1x PBS for 1 h at RT. Primary antibodies were prepared in blocking solution at concentrations detailed in the antibody section. Devices were incubated with primary antibodies for 1-2 h at RT, then washed 3 times with 1x PBS for 10 min each. Secondary antibodies and fluorescent dyes were prepared in blocking solution at concentrations detailed in the antibody section. Devices were incubated with the solution of secondary antibodies and fluorescent dyes for 1-2 h at RT protected from light, then washed 3 times with 1x PBS for 10 min each. Devices were then stored in 1x PBS + 1% penicillin/streptomycin (Sigma) prior to imaging.

#### Confocal microscopy of fixed samples

Confocal microscopy of fixed samples was conducted on two separate setups. The first setup was a Yokogawa CSU10 spinning disk confocal on a Nikon TE2000 microscope with a Cool-SNAP HQ2 camera (Photometrics), controlled by NIS Elements software (Nikon). A Plan Apochromat λ NA 0.75 20x air objective (Nikon) was utilized for both 2D coverslip samples and 3D microvessel samples. Calibration was 0.21 μm/pixel. 405 nm, 488 nm, 561 nm, and 640 nm solid-state lasers were used at 100% power. The second setup was a Yokogawa CSU-W1/SoRa spinning disk confocal with an ORCA Fusion BT sCMOS camera (Hamamatsu) controlled through NIS Elements software (Nikon) (Dema et al., 2024). An Apochromat TIRF 60x NA 1.49 oil immersion objective was utilized for 2D coverslip samples (0.11 μm/pixel), and an Apochromat long working distance 40x NA 1.25 water immersion objective was utilized for 3D microvessel samples (0.16 μm/pixel). 405 nm, 488 nm, 561 nm, and 640 nm solid-state lasers were used at powers between 25-50%.

### Live-cell microscopy

#### Fluorescent 2D live-cell imaging

For live imaging of actin dynamics, either SPY555-FastAct (Spirochrome) was incubated at 1:1000 in medium for 2 h before imaging, or lentiviral LifeAct-mScarlet was transduced into cells 2+ days prior to imaging. For live imaging of NMIIB, lentiviral mEmerald-NMIIB was transduced into cells 2+ days prior to imaging. Cells were imaged on MatTek dishes (poly-D-lysine coated) with an Ibidi 2-well insert for visualization of two cell lines at once. Imaging was conducted on a Yokogawa CSU-X1 spinning disk confocal custom-modified by Spectral Applied Research on a Nikon Ti-E microscope with an Evolve EMCCD camera (Photometrics) controlled by NIS Elements software (Nikon) (Stehbens et al., 2012). A Nikon CFI Apo TIRF NA 1.45 60x oil immersion objective lens was used along with 100-mW diode-pumped solid-state (DPSS) 488 nm (Coherent Sapphire) and 561 nm (Cobolt Jive). Laser power was set to 50% and time-lapse acquisition was taken every 2 or 3 min. Calibration was 0.09 μm/pixel. The imaging system was enclosed within an imaging chamber (In Vivo Scientific) at 37°C with humidified air at 5% CO_2_.

#### Phase 3D live-cell imaging

Microfluidic devices were loaded onto a plastic 6 well plate with PBS-soaked Kimwipes to maintain humidity. The plate was enclosed within an environmental chamber (Okolab) at 37°C with humidified air at 5% CO_2_. Phase imaging was conducted on a Yokogawa CSU10 spinning disk confocal on a Nikon TE2000 microscope with a Cool-SNAP HQ2 camera (Photometrics), controlled by NIS Elements software (Nikon). A Nikon Ph1 ADL NA 0.25 10x phase air objective (0.45 μm/pixel) was utilized and acquisitions were taken every 3 min.

### Traction force microscopy

Compliant polyacrylamide gels were synthesized as in the section “Hydrogel coverslip preparation” but on 25 mm coverslips with 1:100 200 nm green fluorescent beads mixed into the H2O before mixing with other gel ingredients. hMVECs were seeded onto bead gels and reached confluency before imaging. Coverslips were mounted into a live imaging chamber (In Vivo Scientific) at 37°C with humidified air at 5% CO_2_. Imaging was conducted on a Yokogawa CSU-X1 spinning disk confocal custom-modified by Spectral Applied Research on a Nikon Ti-E microscope with an Evolve EMCCD camera (Photometrics) controlled by NIS Elements software (Nikon) (Stehbens et al., 2012). Apochromat 40x NA 1.25 water immersion objective (0.16 μm/pixel) was used along with 100-mW diode-pumped solid-state (DPSS) 488 nm (Coherent Sapphire) and 561 nm (Cobolt Jive). Images of beads were acquired before and after detachment of cells with 10% sodium dodecyl sulfate. Traction force vector fields were created by comparing the tensed (pre-detachment) to relaxed (post-detachment) images of beads using open-source FIJI packages for image registration, iterative particle image velocimetry, and Fourier transform traction cytometry (Tseng et al., 2012), approximating the gel stiffness as 2.5kPa and regularization factor as 10^-10^.

### Image processing and analysis

For all quantification of immunofluorescence images, regions of interest with large holes in the monolayer were excluded. Within an independent experiment, only regions of interest with similar cell density were compared across conditions. Unless otherwise mentioned, all representative images are intensity-matched across conditions for data visualization.

#### AJ morphology

AJ morphology was assessed using JunctionMapper software (Brezovjakova et al., 2019). For each image, a junctional skeleton mask was created based on maximum intensity of the junction stain. Operations were conducted on each junction individually. For junction linearity, the contour of each junction was compared to the straight-line distance between the two ends of the junction: closer to 1 indicates a linear junction, while greater than 1 indicates a more undulated/wavy junction. To obtain junction width, the junction skeleton mask was dilated 10x from the cell border. Fluorescence above a set threshold for each biological replicate was recognized as junctional signal. From this signal, junctional area was quantified. Junction width was calculated as junctional area divided by junction contour length. Outliers were removed in Prism using ROUT method (Q = 0.1%). For each biological replicate, individual values were normalized to the mean Scramble value.

#### Junctional intensity

Junctional intensity of immunostained VE-cadherin and catenin proteins was analyzed using ImageJ. Single planes of confocal images were cropped into patches of 100-200 μm depending on the field of view offered by the microscope. VE-cadherin staining was thresholded at a constant value for each biological replicate and used to create a junctional mask. Mean fluorescence intensity was measured within this mask. For each biological replicate, individual values were normalized to the mean Scramble value.

#### Actomyosin distribution – cortical:interior ratio

Single planes of confocal images with the actomyosin marker of interest were analyzed in ImageJ. For each image, a junctional skeleton mask was created using JunctionMapper software (Brezovjakova et al., 2019). This mask was dilated in ImageJ 10x to generate the “cortical” area of the image, while the inverse was the cell “interior”. For each field of view, we quantified mean fluorescence intensity of actomyosin staining within the mask (“cortical”) relative to mean fluorescence intensity of actomyosin staining outside the mask (cell “interior”). A ratio greater than 1 indicates higher abundance of actomyosin in the junctional cortex, where a ratio near 1 indicates that the abundance of actomyosin in the junctional cortex is similar to the abundance of actomyosin in the cell interior.

#### Actomyosin distribution – line scans

Single planes of confocal images with the actomyosin marker of interest were analyzed in ImageJ. Using a junctional co-stain, a line scan was drawn from cell edge to cell edge through the short axis of the nucleus and the brightest regions of the actomyosin cortex at the edges. Fluorescence intensity along this line was quantified via line scan in ImageJ and normalized to the max intensity observed in Scramble for that biological replicate to achieve values between zero and 1 (a.u.). For each cell, width was normalized to the maximum x position to achieve values between zero and 1 (a.u.). Cells of different sizes were compared by interpolation in MATLAB with a spacing of 0.001 from x=0 to x=1, where x=1 is the normalized width of cells measured. Finally, each cell was split into halves and analyzed from x=0 (cell periphery, VE-cadherin border) to x=0.5 (cell center). To highlight the “cortical” localization for this distribution (between x=0 and x=0.25), the line scan fluorescence intensity values were normalized to average intensity of cell interior of each condition (x=0.25 to x=0.5).

#### Representative line scans

Single planes of confocal images were analyzed in ImageJ. A line scan was drawn over the feature of interest, and fluorescence intensity was measured for each channel/stain. For each channel’s line trace, fluorescence intensity values were normalized to the minimum and maximum fluorescence values. The line distance was normalized by setting the maximum distance to 1. Traces were plotted in Prism using 6^th^ order smoothing.

#### Scrib cluster area

Confocal images of Scrib immunostaining were used to create maximum intensity projections in ImageJ. Images were cropped into patches of 100-200 μm depending on the field of view offered by the microscope. The patch was duplicated and thresholded with a constant value for each biological replicate that eliminated background Scrib signal. This threshold was applied to create a mask, and the watershed function was applied 4 times to separate grouped clusters. “Analyze Particles” was used; inclusion criteria for cluster area were determined by comparing Scramble to *SCRIB^KO^* and setting a cutoff that excluded most *SCRIB^KO^* signal (0.3 μm^2^ and above). Particles were sent to ROI manager, a selection was created, and moved to the original image. Each particle area was quantified using a macro to iterate measurements across all ROI. Outliers were removed in Prism using ROUT method (Q = 0.1%).

#### Sprout metrics

Sprout metrics were quantified using phase images of full-length vessels stitched in ImageJ (Preibisch et al., 2009). For each vessel, a set length of 2 mm or 3 mm in the center of the 4 mm parent vessel length was used for quantification to exclude aberrations from the vessel entry site into the ECM chamber. Using this quantification length, sprouting events (intact sprouts and detached cells) were calculated and reported as feature per mm of parent vessel. Sprout length was taken as the longest intact sprout from each parent vessel within this quantification area. Inevitably, several vessels from each condition in each experiment had a few sprouting events pre-treatment. To account for this baseline, each parent vessel was normalized to itself by subtracting the number of sprouts or detached cells pre-treatment from the post-treatment values. Sprout length was normalized internally to each parent vessel by subtracting the value for sprout length pre-treatment. Detached cells that were present on day 0 but not visible on day 1 (migrated out of field of view or apoptosed) were accounted as part of the detached cell count on day 1 so that “negative” detached cell values were not obtained. In fixed confocal images of phalloidin/DAPI-stained microvessels from microvessels fixed at 24h or 72h S1P treatment, max projections of half the Z-stack of parent vessels were used for quantification of sprout parameters and representative images. Inclusion criteria for sprout parameters were intact sprouts that were not obscured by another sprout, with surface clearly visible and not obscured by high intensity of vessel edge. An ROI of 50 μm wide x 25 μm tall spanning the sprout – parent vessel interface (“sprout base”) was used for quantification of actin cluster counts within this interface. The broadest point of the sprout base was measured for sprout base width.

### Statistical analysis

All statistical analysis was conducted in GraphPad Prism 10. Data were tested for normal distribution via D’Agostino & Pearson test or Shapiro-Wilk test (alpha = 0.05) depending on sample size. For comparisons between two conditions, normally distributed data were analyzed via unpaired two-tailed student’s t-test, and non-normally distributed data were analyzed via Mann-Whitney U test. For comparisons between more than two conditions, normally distributed data were analyzed via two-way ANOVA with Tukey’s multiple comparisons test, and non-normally distributed data were analyzed via Kruskal-Wallis test with Dunn’s multiple comparisons test. “rel.” indicates that data points (n) were normalized to the mean control value for each independent biological replicate (N). Bar graphs presented mean +/- standard deviation. Data analysis was not blinded and experiments were not randomized.

## SUPPLEMENTAL INFORMATION

**Supplementary Figure 1:**
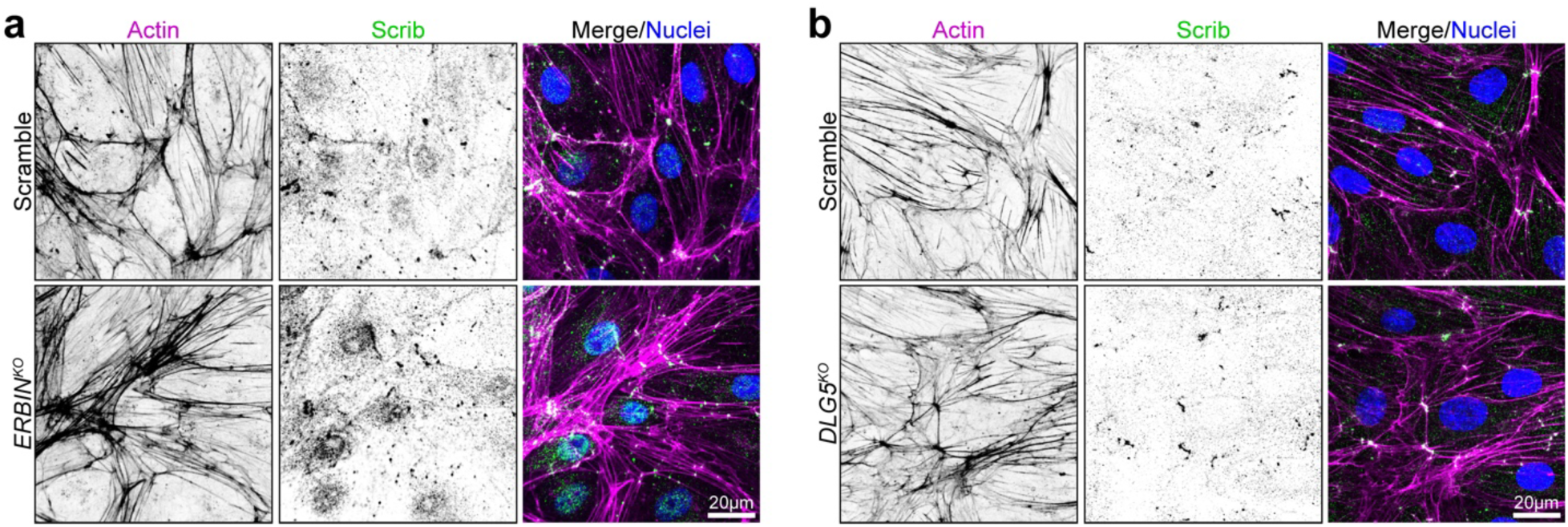
Scrib localization is unchanged in *ERBIN^KO^* and *DLG5^KO^*. Representative confocal micrographs of actin and Scrib in hMVECs, comparing **(a)** Scramble and *ERBIN^KO^* (N = 1 independent experiment) and **(b)** Scramble and *DLG5^KO^* (N = 1 independent experiment).

**Supplementary Figure 2:**
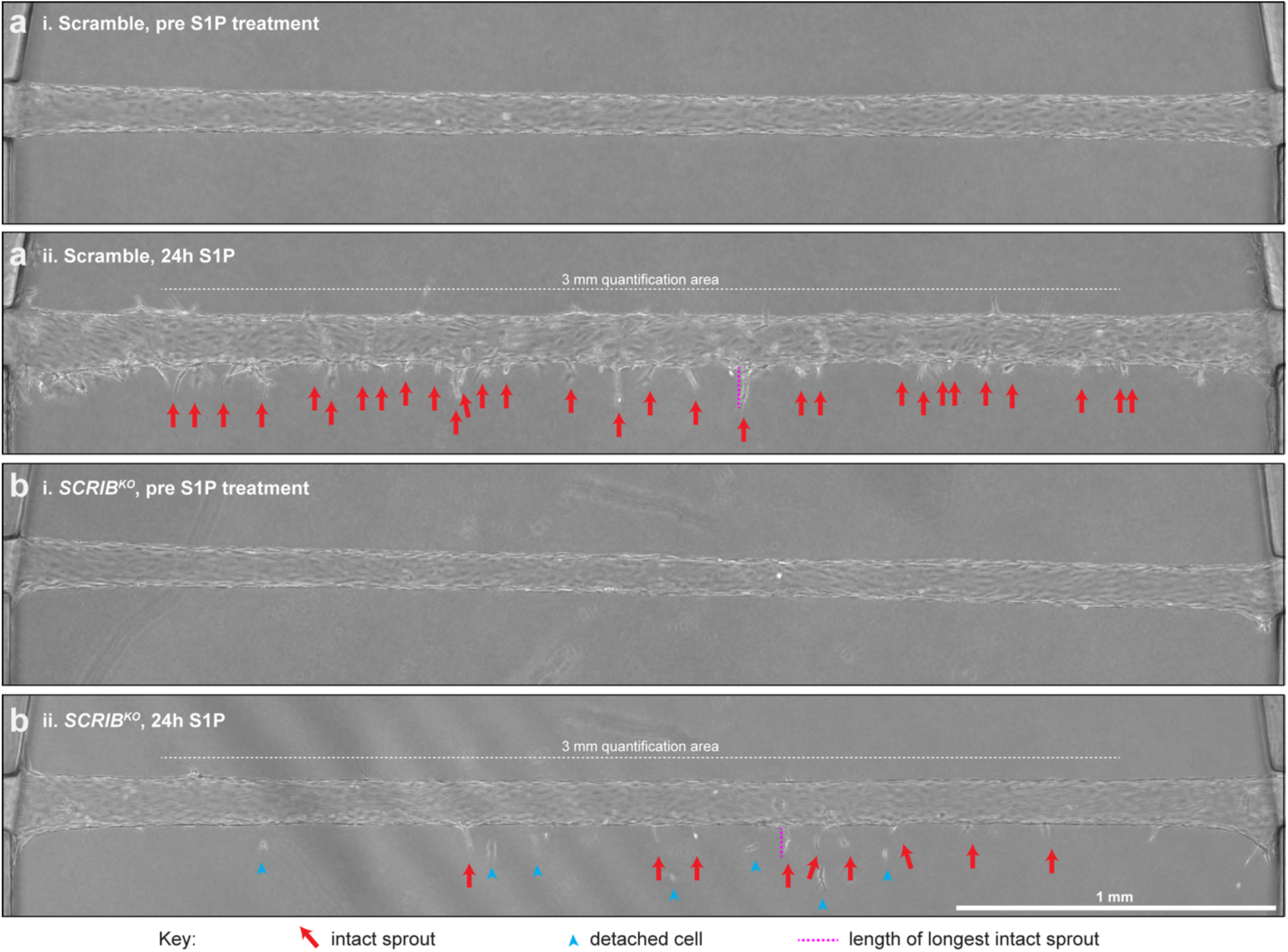
Full-length Scramble and *SCRIB^KO^* microvessels and quantification parameters. Representative full-length images of **(a)** Scramble and **(b)** *SCRIB^KO^* hMVEC microvessels before (i) and after (ii) 24 h 500 μM S1P treatment (N = 3 independent experiments, n = 3-5 devices per condition per N). All parameters at 24 h are quantified over a set distance, in this case 3 mm, and reported per mm. Red arrows indicate intact sprouts, blue arrowheads indicate detached cells, and dotted pink lines indicate the length of the longest intact sprout per vessel.

**Supplementary Figure 3:**
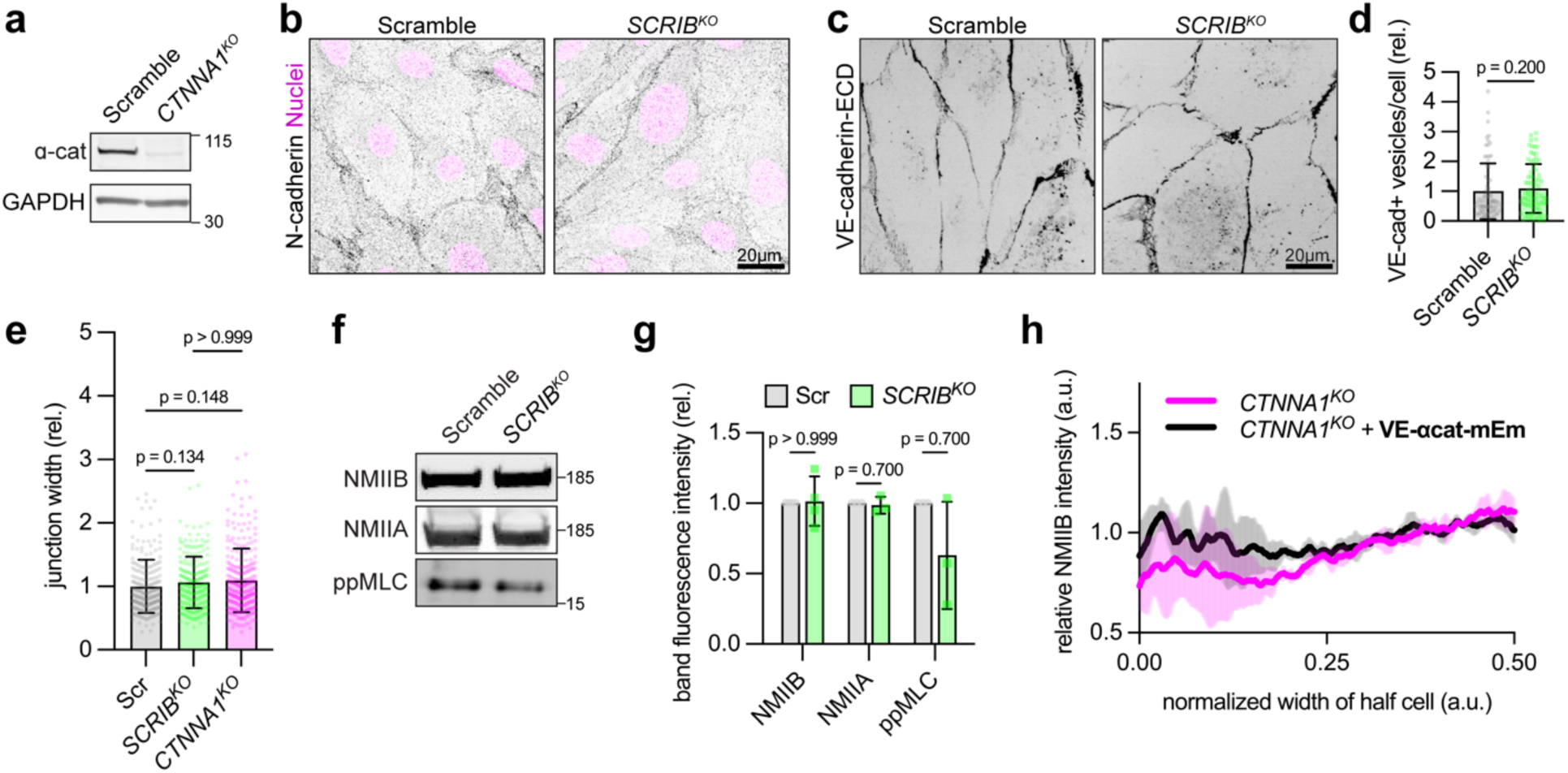
*SCRIB^KO^* impact on adherens junctions and associated actomyosin. **(a)** Lysates from Scramble and *CTNNA1^KO^* blotted for ⍺-catenin (N = 1). **(b)** Immunofluorescence staining of N-cadherin in Scramble and *SCRIB^KO^* (N = 3 independent experiments). **(c)** Visualization of VE-cadherin internalization through immunofluorescence staining of antibody targeting the extracellular domain of VE-cadherin after live antibody incubation. **(d)** Quantification of count of VE-cadherin+ vesicles per cell in Scramble and *SCRIB^KO^* (N = 3 independent experiments, n ≥ 20 cells per condition per N, Mann-Whitney test). **(e)** Quantification of VE-cadherin junction width in Scramble, *SCRIB^KO^*, and *CTNNA1^KO^* (N = 4 independent experiments, n ≥ 40 junctions per condition per N, Mann-Whitney test). **(f)** Lysates from Scramble and *SCRIB^KO^*cells blotted for NMIIB, NMIIA, and ppMLC (Thr18/Ser19). **(g)** Quantification of Western blot band fluorescence intensities from (f) (N ≥ 3, multiple Mann-Whitney tests). **(h)** Normalized NMIIB cortical abundance in *CTNNA1^KO^*, with or without VE-αcat-mEm expression (N = 3 independent experiments, n ≥ 20 cells per condition per N).

**Supplementary Figure 4:**
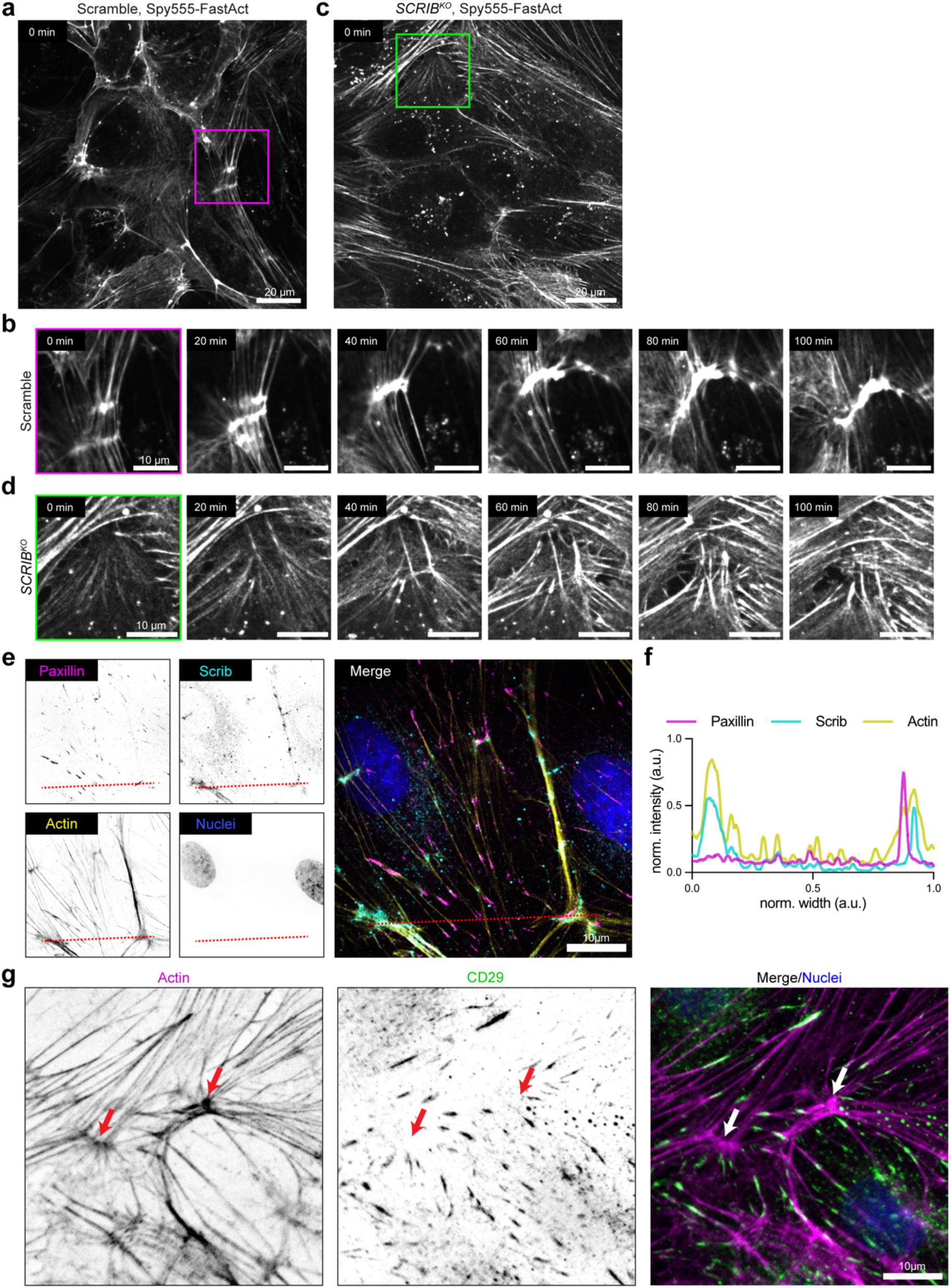
Actomyosin clusters form and coalesce in Scramble but fail to do so in *SCRIB^KO^*; Scrib/actomyosin clusters do not overlap focal adhesion components. **(a, b)** Representative live actin imaging (Spy555-FastAct) in Scramble hMVEC monolayer. Inset highlights clusters that coalesce into a larger cluster over 100 min (Video 3) N = 3 independent experiments). **(c, d)** Representative live actin imaging (Spy555-FastAct) in *SCRIB^KO^* hMVEC monolayer. Inset highlights actin fibers that coalesce but ultimately fail to form a large cluster over 100 min (Video 4) (N = 3 independent experiments). **(e)** Representative immunofluorescence staining of Scrib in hMVECs co-stained with Paxillin and actin (N = 2 independent experiments). **(f)** Line scan of representing the three markers along the line demarcated in (e). **(g)** Representative immunofluorescence staining of activated β1 integrin (CD29) co-stained with actin (actin clusters, arrows) (N=3 independent experiments).

**Supplementary Figure 5:**
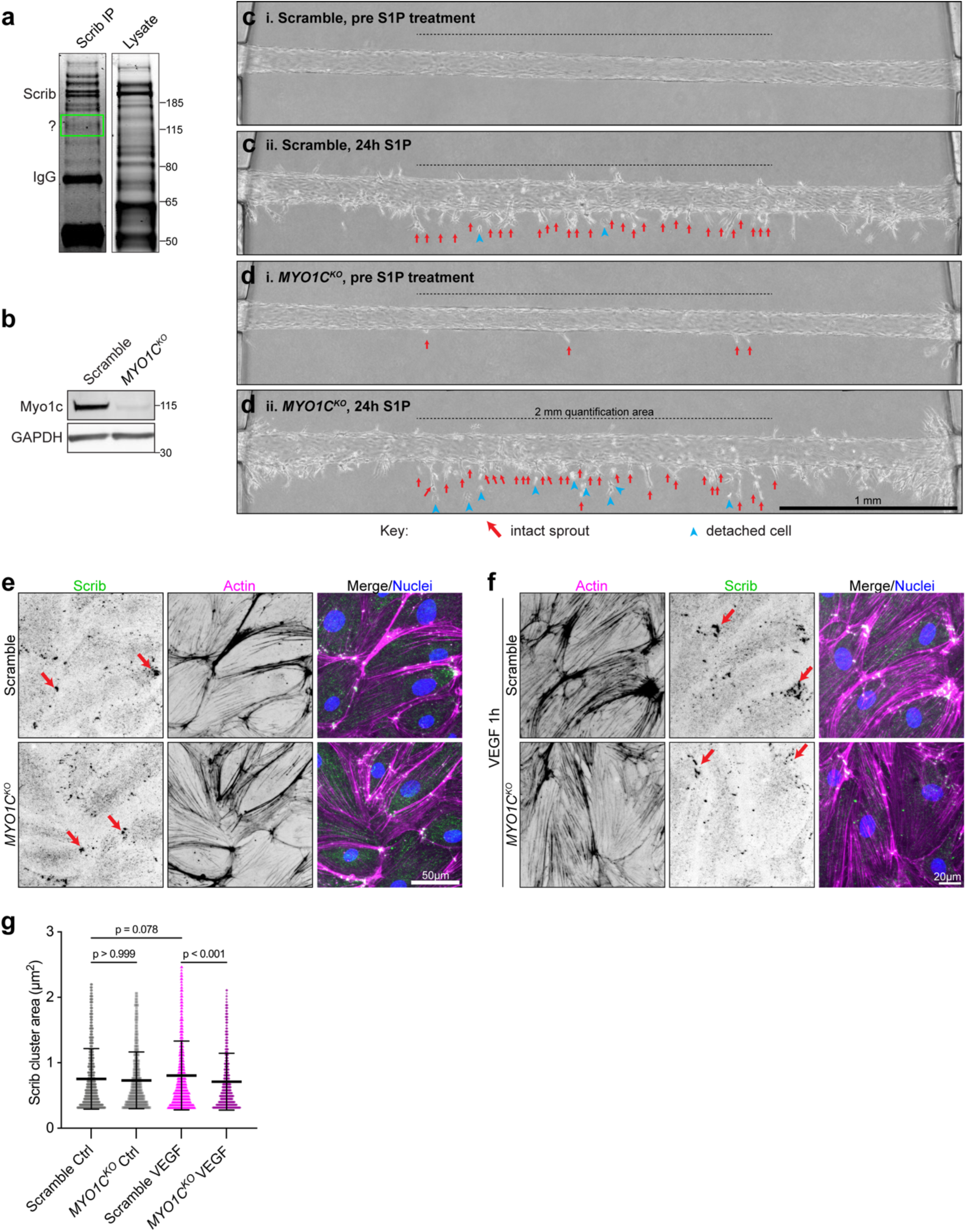
Myo1c regulates nascent sprout integrity and cluster size in the angiogenic context. **(a)** Wild-type hMVEC lysate immunoprecipitated with anti-Scrib antibody. Immunoprecipitate and whole lysate were visualized by Coomassie Blue stain. A band was excised at ∼120 kDa and sent to mass spectrometry for protein identification (N = 1 for mass spectrometry, observed prominent band in Coomassie N = 4 times). **(b)** Immunoblot of Myo1c in Scramble and *MYO1C^KO^* (N = 2). **(c, d)** Representative full-length images of **(c)** Scramble and **(d)** *MYO1C^KO^* hMVEC microvessels before (i) and after (ii) 24 h 500 uM S1P treatment (N = 4 independent experiments, n ≥ 3 devices per condition per N). All parameters at 24h are quantified over a set distance, in this case 2 mm, and reported per mm. Red arrows indicate intact sprouts; blue arrowheads indicate detached cells. **(e-f)** Representative immunostaining of Scrib clusters and actin within hMVEC monolayers, comparing untreated cells to cells stimulated by VEGF (50 ng/ml, 1 hr) for both Scramble *MYO1C^KO^*. **(g)** Quantification of Scrib cluster size (N = 3 independent experiments, n ≥ 400 clusters per condition per N, Mann-Whitney test).

**Video 1: Live sprouting from parent microvessel captured with phase imaging, Scramble vs. *SCRIB^KO^* (Fig. 2i).** hMVECs composing a 3D microvessel in angiogenesis model after treatment with 500 μM S1P for 12 h, Scramble vs. *SCRIB^KO^*, visualized via time-lapse phase microscopy. Frames were collected every 3 min. Time scale in hh:mm and displayed at 10 frames per sec. Corresponds to Figure 2i.

**Video 2: Spy555-FastAct imaging of confluent endothelial monolayer highlighting cluster formation during retrograde actin flow from cell-cell adhesion, Scramble vs. *SCRIB^KO^* (Fig. 5c).** hMVECs pre-labeled with Spy555-FastAct for 2 h, Scramble vs. *SCRIB^KO^*. Visualized via time-lapse confocal microscopy. Frames collected every 2 min. Time scale in min and displayed at 5 frames per sec. Corresponds to Figure 5c.

**Video 3: Actin clusters in Scramble often coalesce into larger clusters (Fig. S4a, b).** Scramble hMVECs pre-labeled with Spy555-FastAct for 2 h. Visualized via time-lapse confocal microscopy. Frames collected every 2 min. Time scale in min and displayed at 5 frames per sec. Corresponds to Supplementary Figure 4a, b.

**Video 4: *SCRIB^KO^* actin occasionally forms cross-linked arrayed structures but fails to coalesce into clusters (Fig. S4c, d).** *SCRIB^KO^* hMVECs pre-labeled with Spy555-FastAct for 2 h. Visualized via time-lapse confocal microscopy. Frames collected every 2 min. Time scale in min and displayed at 5 frames per sec. Corresponds to Supplementary Figure 4c, d.

**Video 5: Dynamic clustering behavior of non-muscle myosin IIB occurs in Scramble but not *SCRIB^KO^* (Fig. 6b).** hMVECs labeled via lentiviral-mediated expression of mEmerald-NMIIB, Scramble vs. *SCRIB^KO^*. Visualized via time-lapse confocal microscopy. Frames collected every 3 min. Time scale in min and displayed at 5 frames per sec. Corresponds to Figure 6b.

**Supplementary Table 1:** Primers for generating CRISPR KO plasmids.

**Supplementary Table 2:** Primers for DNA fragment assembly.

**Supplementary Table 3:** VE-cadherin BioID – list of proteins identified by mass spectrometry.

**Supplementary Table 4:** Abbreviations.

**Source Data 1:** Full blots from all figures.

**Source Data 2:** Source data for graphs from all figures.

