## Supplementary material for "Scrib organizes cortical actomyosin clusters to maintain adherens junctions and angiogenic sprouting": Source Data 1: Full Blots

Figure 1c

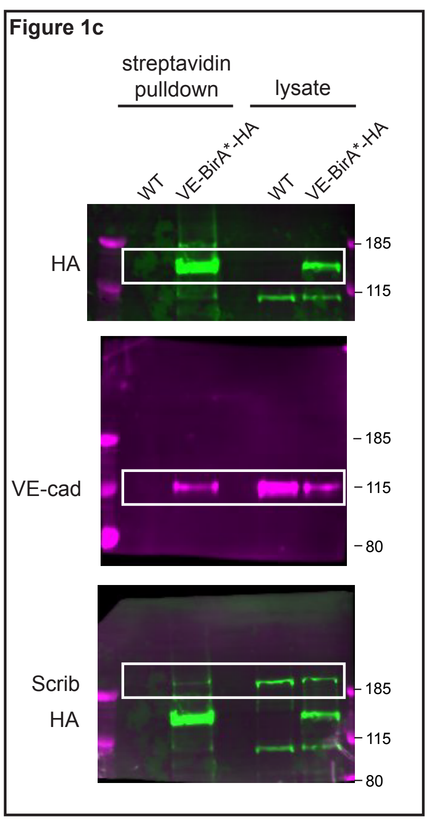


Figure 1d

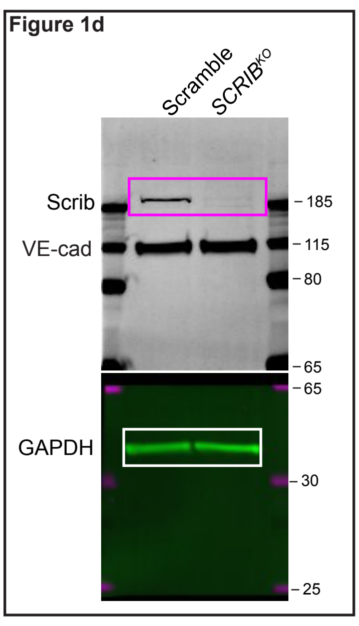


Figure S3a

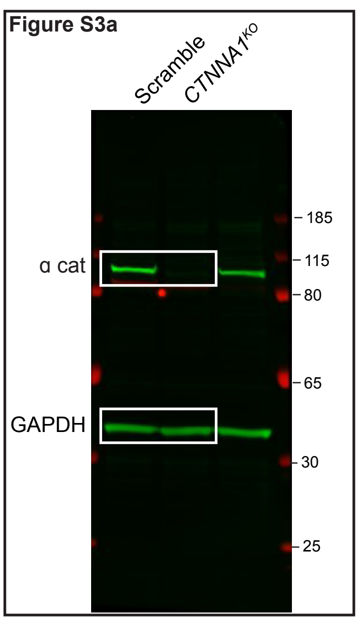


Figure 3g

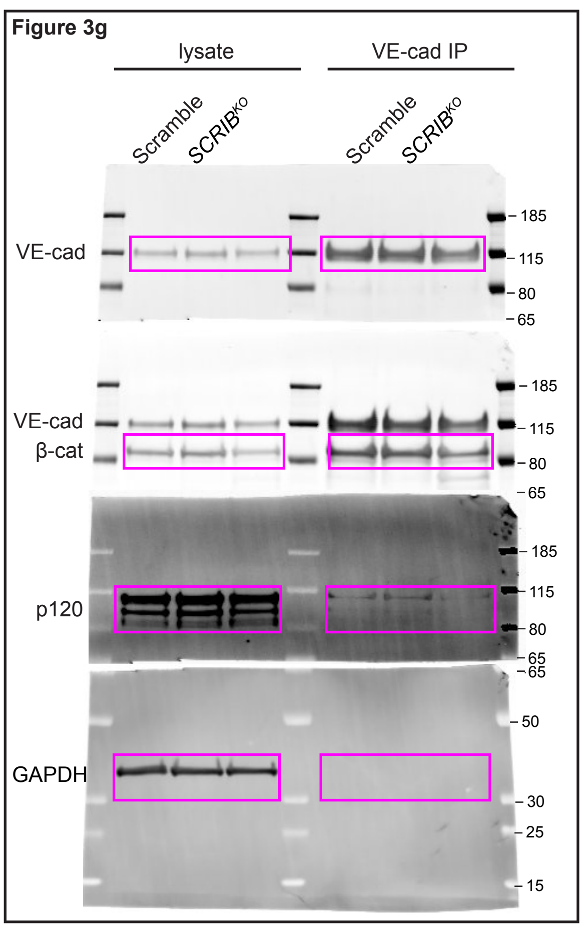


Figure S3f

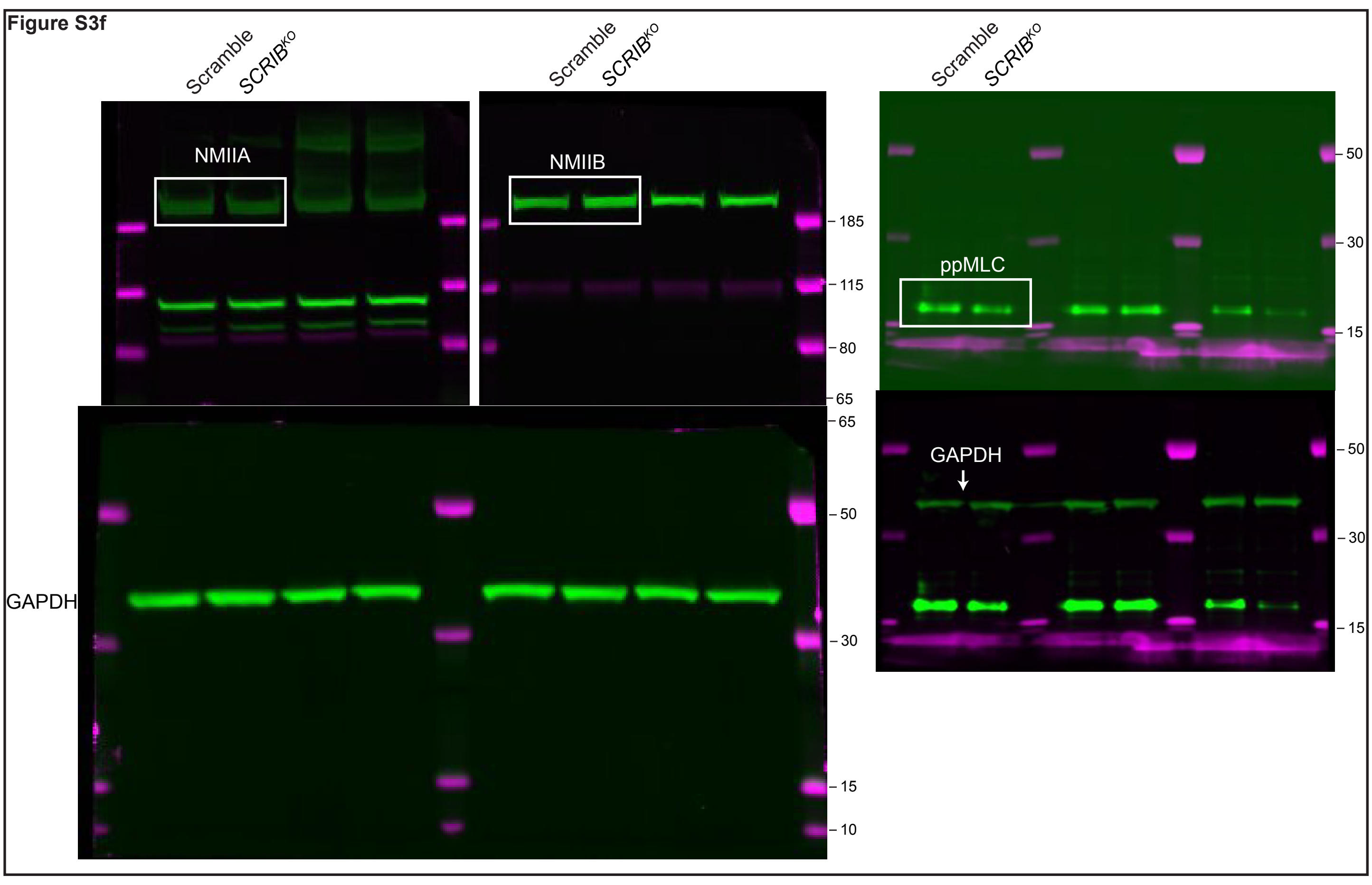


Figure 7a

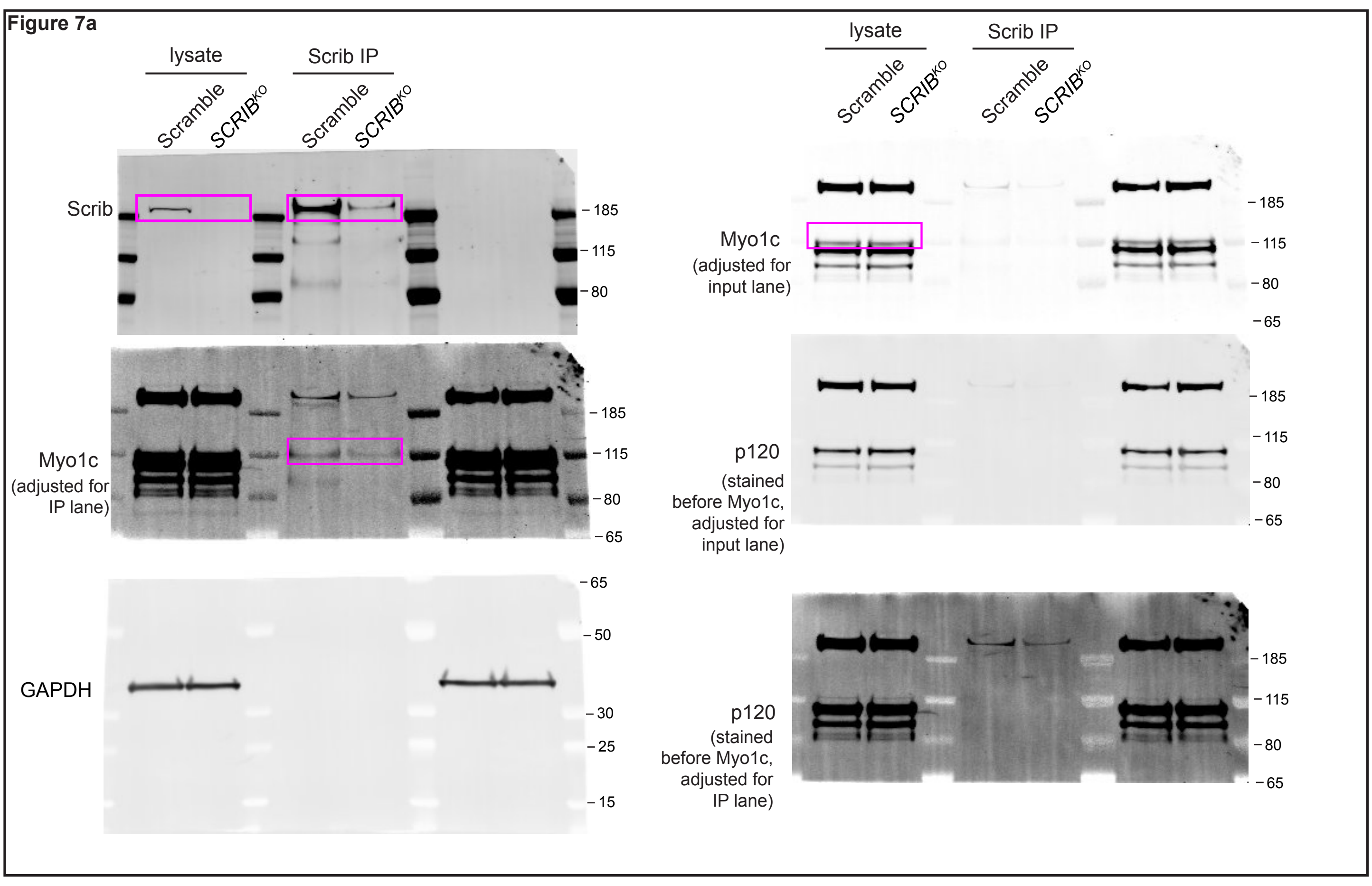


Figure 7b

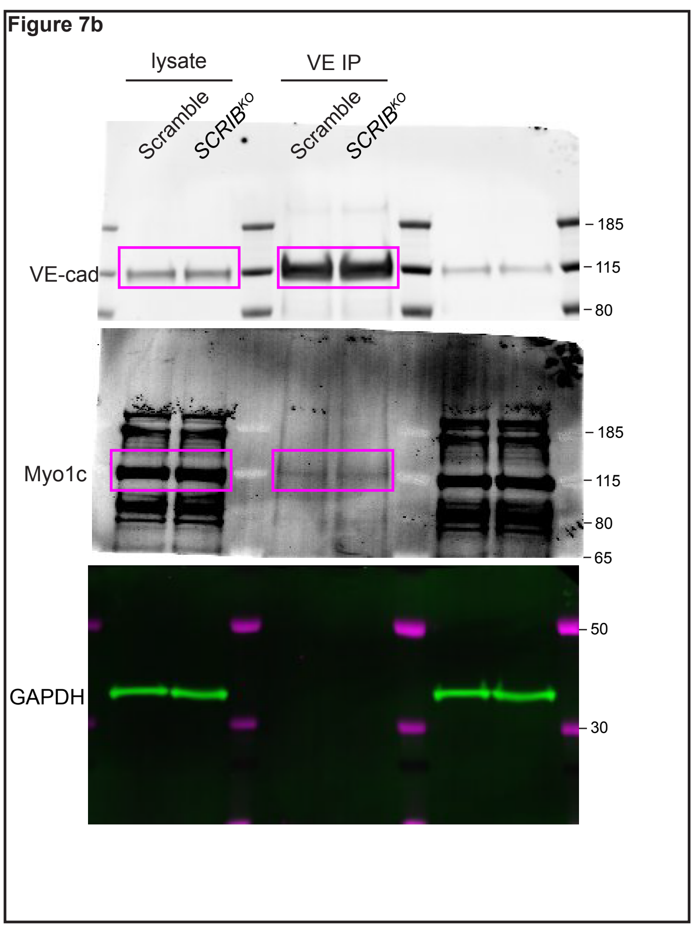


Figure S5a Coomassie blue stain (same gel, brightness adjusted for each)

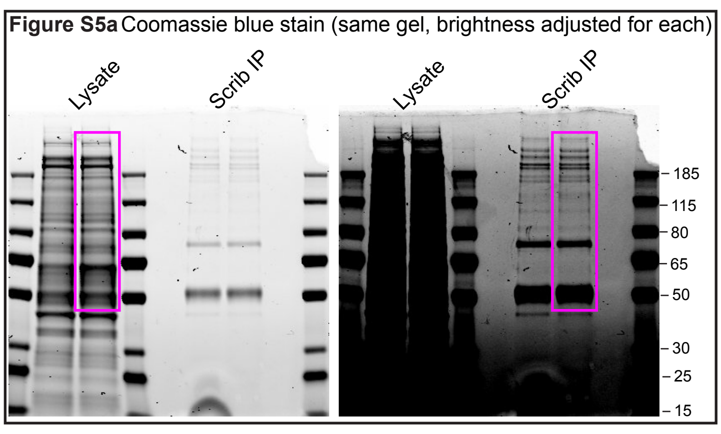


Figure S5b

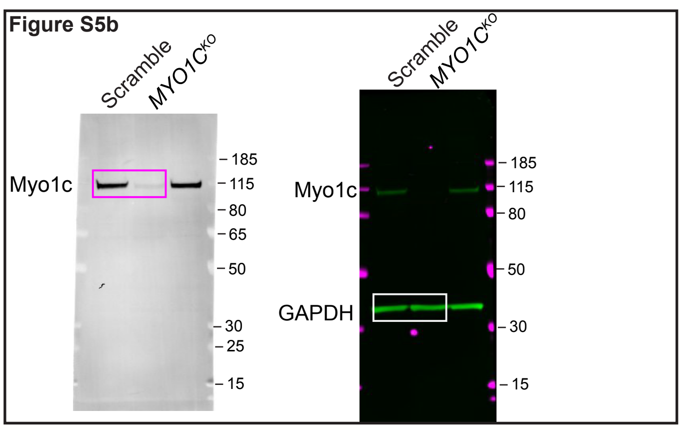
